# Alzheimer’s disease risk gene *Wwox* protects against amyloid pathology through metabolic reprogramming

**DOI:** 10.1101/2025.05.01.651195

**Authors:** Hannah J Lucas-Clarke, Daniel C Maddison, Leonardo Amadio, Edgar Buhl, Katherine O’Hare, Clement Regnault, Owen M Peters, James J L Hodge, Gaynor A Smith

## Abstract

Genome wide association studies have identified multiple loci that mediate the risk of developing late-onset Alzheimer’s Disease (LOAD). The gene WW-domain containing oxidoreductase (*WWOX*) has been identified in recent LOAD risk meta-analyses, yet its function in the brain is poorly understood. Using *Drosophila,* we discovered that knockdown of the highly conserved *Wwox* gene impacts longevity and sleep, having roles in both neuronal and glial subtypes. In an amyloid beta 42 (Aβ_42_) transgenic model of AD, RNAi-mediated knockdown of *Wwox* significantly decreased both lifespan and locomotion whilst elevating soluble Aβ_42_. Transcriptomic and metabolomic analyses revealed that these effects were accompanied by elevated lactate dehydrogenase (*Ldh*) mRNA and lactate levels, downstream of an increase in the key unfolded protein response protein Atf4. Strikingly, we found that upregulation of *Wwox* in the Aβ_42_ model through CRISPR activation significantly reduced amyloid load, improved longevity and locomotion. Multi-omics analysis revealed *Wwox* upregulation partially reversed several key Aβ_42_-induced transcriptional pathways in the brain and reduced levels of L-methionine and associated enzymes. These findings support a role for reduced WWOX levels in the genetic risk of developing LOAD via pyruvate metabolism and point towards WWOX activation as a protective therapeutic strategy.

## Introduction

Alzheimer’s Disease (AD) is the most common form of dementia, characterised by pronounced neurodegeneration, causing memory impairment, cognitive decline, and sleep disruption. On a cellular and molecular level, these symptoms follow the accumulation of extracellular amyloid β (Aβ) plaques, hyperphosphorylated tau pathology and reactivity of astrocytes and microglia. AD manifests in two forms, early-(EOAD) and late-onset (LOAD). Whilst EOAD can be caused by monogenic mutations in *amyloid precursor protein (APP), presenilin-1 (PSEN1*) or *presenilin-2 (PSEN2*), LOAD is considered sporadic and a result of the convergence of environmental and genetic risk factors. Single nucleotide polymorphisms (SNPs) at approximately 90 genomic loci have been discovered to contribute to LOAD risk (1). Determining how genes associated with these SNPs contribute to disease onset and progression is vital to our understanding of LOAD and how it may be prevented or treated.

An intergenic SNP located on chromosome 16 between *WW-domain containing oxidoreductase* (*WWOX*) and *MAF bZIP transcription factor* (*MAF*) has been associated with an increased the risk of developing LOAD (2). Genome-wide expression quantitative trait locus (eQTL) mapping between LOAD patients and controls identified *WWOX* as a significant eQTL in the brain with a negative β value, indicating a decrease in *WWOX* mRNA in individuals with the associated SNP (3). A genome-wide significant association has also been made between a SNP within a predicted *WWOX* enhancer region and a key disease biomarker, Aβ_42_/Aβ_40_ ratio (4). Furthermore, decreases in WWOX have been observed in post-mortem hippocampal tissue of AD patients compared to controls (5). However, the mechanisms underlying *WWOX* modulation of AD are not well understood.

WWOX is an oxidoreductase and tumour suppressor gene that plays a significant role in cancer cell metabolism, influencing glycolysis, mitochondrial integrity, signalling and oxidative stress (6–10). In addition, *WWOX* loss of function has been postulated to mediate a Warburg-like effect – a switch to glycolysis despite aerobic conditions, via modulation of HIF1α, a transcription factor with a variety of metabolic targets including *glucose transporter 1* (*GLUT1*) and *lactate dehydrogenase* (*LDH*) (6,11,12). A similar metabolic shift has been observed in *Drosophila* (13) and human stem cell-derived neuronal models of AD (14,15), but whether this is influenced by WWOX, in the context of AD, has not yet been explored.We sought to address this question by altering *Wwox* levels in the *Drosophila* brain, via neuronal or glial-specific RNA interference (RNAi) and CRISPR activation (CRISPRa). We explored the age associated effects on AD-relevant phenotypes with or without the overexpression of the human Aβ_42_ peptide (16). We show that both neuronal and glial knockdown of *Wwox* shortened lifespan and disrupted sleep in flies, in the absence of Aß_42_. However, in the presence of Aβ_42,_ *Wwox* knockdown exacerbates lifespan and locomotor deficiencies in a neuron-specific manner, whilst also increasing amyloid load. Transcriptomic and metabolomic analyses identified that elevated *Ldh*, as a consequence of Aβ_42_ overexpression, was exacerbated by *Wwox* knockdown and was accompanied by increased levels of lactate and the integrated stress response (ISR) / unfolded protein response (UPR) regulator Atf4. Strikingly, upregulation of *Wwox* in neurons significantly reduced amyloid load, rescuing lifespan and locomotor defects but did not alter lactate levels, instead lowering L-methionine. Taken together, these findings provide insight into the mechanisms underlying LOAD risk conferred by genetic variation at the *WWOX* locus and identify WWOX activation as a strategy for reducing Aβ_42_ pathology.

We sought to address this question by altering *Wwox* levels in the *Drosophila* brain, via neuronal or glial-specific RNA interference (RNAi) and CRISPR activation (CRISPRa). We explored the age associated effects on AD-relevant phenotypes with or without the overexpression of the human Aβ_42_ peptide (16). We show that in the absence of Aß_42_ both neuronal and glial knockdown of *Wwox* shortened lifespan and disrupted sleep. However, in the presence of Aβ_42,_ *Wwox* knockdown exacerbates lifespan and locomotor deficiencies in a neuron-specific manner, whilst also increasing amyloid load. Transcriptomic and metabolomic analyses identified that elevated *Ldh*, as a consequence of Aβ_42_ overexpression, was exacerbated by *Wwox* knockdown and was accompanied by increased levels of lactate and the integrated stress response (ISR) / unfolded protein response (UPR) regulator Atf4. Strikingly, upregulation of *Wwox* in neurons significantly reduced amyloid load, rescuing lifespan and locomotor defects but did not alter lactate levels, instead lowering L-methionine. Taken together, these findings provide insight into the mechanisms underlying LOAD risk conferred by genetic variation at the *WWOX* locus and identify WWOX activation as a strategy for reducing Aβ_42_ pathology.

## Materials and Methods

### Fly husbandry

*Drosophila* stocks (Table.S1) were housed at room temperature and transferred to fresh standard ‘Cornmeal-Molasses-Yeast’ media, set with agar once every 3 weeks. Crosses and experimental lines were housed at 25 °C on a 12:12 hour light-dark schedule.

### Single cell atlas of the ageing Drosophila brain

*Wwox* expression was assessed using a single cell transcriptomic dataset of the ageing *Drosophila* brain available on SCOPE (17). We calculated the percentage of cells within annotated clusters that expressed *Wwox*.

### Lifespan

Newly eclosed females were allowed to mate for <48 hours, then were separated into vials of n<11 and passed to fresh food 2-3 times a week, whilst scoring deaths. Alive flies that were not transferred successfully into the new vial were censored in the analysis. Kaplan-Meier plots were generated (GraphPad Prism v.10) and analysed using Mantel-Cox and Cox proportional hazard-model tests, correcting for multiple comparisons using 5 % FDR, where necessary.

### Rapid Iterative Negative Geotaxis (RING)

Newly eclosed females were allowed to mate for <48 hours and then divided into vials of n<11. Progeny were flipped onto fresh food 2-3 times a week and assayed at 7-, 14-, 21- and 28-days post eclosion (±1 day). After 5 minutes of acclimatisation, the apparatus was raised to a fixed height (8 cm) and dropped, images were captured once a second for 10 seconds. After 1-minute recovery, this procedure was repeated for a total of five drops. Using FIJI, the mean distance climbed by each fly at the 4 second time point was measured. Data was analysed by repeated measures mixed effect general linear models (genotype x age, matching by vial). If repeated measures were not applicable, ordinary two-way ANOVAs or one-way ANOVAs were performed (GraphPad Prism v.10). For the comparison of two groups, unpaired two tailed t-tests were performed. Significant effects resulting from ANOVA’s were explored further using Šidák multiple comparisons.

### Assessment of sleep and circadian rhythms

Males (0-7 days) were individually housed within *Drosophila* activity monitors (DAM, Trikinetics Inc) and subjected to a 12-hour light:dark (LD) cycle at 25 °C for 5 days, followed by 5 days of 24-hour darkness (DD). For each run (>2 runs), N=31-80 individual males were assayed per genotype. Activity (beam crosses) was calculated in 1- and 30-minute bins using DAMFileScan v1.11 and an in-house script (https://github.com/Fly-Cardiff/sleep_analysis) and subsequently analysed using “Sleep and Circadian Analysis MATLAB Program - SCAMP”(18,19) (MATLAB, vR2020b). A bout of sleep was defined, as per convention, as more than five minutes without activity (20,21). Data from the LD phase was derived after exclusion of day 1, to allow for entrainment, and after removal of any deaths.

Seventeen metrics of sleep and wakefulness were exported from SCAMP for subsequent analyses: total wake episodes, mean wake episode duration, total activity counts, total time awake, latency to sleep, mean wake duration minus latency, mean activity a minute whilst moving, activity normalised to time awake, the probability of transitioning to an inactive state or active state, the number of sleep episodes, the total, max, mean and median sleep episode duration and sleep stability. All metrics were assessed for inter-correlations (Pearsons) and strongly correlated (r>|0.8|) metrics were excluded from further analysis. Of the remaining metrics, normality of residuals was assessed with Kolmogorov-Smirnov tests and quantile-quantile plots and consequent Box-Cox transformations were performed where data was deemed sufficiently non-Gaussian (RStudio v2022.12, GraphPad Prism v.10). Multiple two-way ANOVAs were then performed on the surviving metrics, corrected by 5% FDR. If a significant interaction was identified, Bonferroni multiple comparison tests were performed for pairwise comparisons of interest (GraphPad Prism v.10) and further corrected for multiple testing across metrics.

Assessment of circadian strength was performed for the DD regime, including flies that survive to day 10. Rhythmicity statistic (RS) and period length were derived from SCAMP autocorrelograms. The RS describes the ability of a fly to maintain a cycle, with an RS>1.5 being rhythmic, RS=1-1.5 being weakly rhythmic and flies with an RS< 1 being arrhythmic. Periodicity was calculated for rhythmic and weakly rhythmic flies only. If a fly had a RS<0, it was set to 0 for analysis of rhythmicity. Data was analysed via t-test or non-parametric Mann-Whitney tests dependent on normality (GraphPad Prism v.10).

### Immunohistochemistry

*Drosophila* heads were dissected and fixed in 4 % PFA in PBS with 0.1 % Triton X-100 (PTX) for 10 minutes. Brains were dissected and fixed for a further 10 minutes. Brains were blocked in 10 % normal goat serum (in PTX) for 1 hour at room temperature, then incubated with primary antibodies – anti-Aβ (1-16, 6E10, mouse, 1:400, Biolegend #803001), anti-Atf4 (rat, 1:200, gifted by Bateman lab (22)) – for 60 and 16 hours respectively at 4 °C in 10 % normal goat serum (in PTX). Brains were washed 5 times in PTX, then incubated with fluorophore-conjugated secondary antibodies AlexaFluor-647 (anti-mouse, 1:500, Invitrogen #A-21235), AlexaFluor-594 (anti-rat, 1:400) for 1-2 hours at room temperature in PTX. Brains were then washed 5 times in PTX, then mounted in VECTASHEILD Antifade (Vector Laboratories, H-1000-10) before imaging on a Cell Observer Spinning Disk Confocal Microscope (ZEISS) with Plan Apo 20x/0.8 dry objective and Axiocam 503 monochromatic camera, using Zen Blue software (ZEISS). Images were processed in FIJI - Aβ aggregates were detected by the *Analyze Particles* function and Atf4 fluorescence intensity (mean gray value) was averaged across 4 equal sized ROIs (227.5 μm^2^) per brain. Aβ intensity was analyzed with one-way ANOVA or t-test depending on the number of groups, Atf4 fluorescence was assessed via Mann-Whitney tests (GraphPad Prism v.10).

### Imaging of sima:GFP fluorescent reporter

Brain sima levels were measured using the genetically encoded reporter line UAS-sima:GFP. Unfixed brains were dissected and imaged on a Zeiss Cell Observer spinning disk confocal using 488nm excitation, Plan Apo 20x/0.8 dry objective and Axiocam 503 camera. Fluorescence intensity (mean gray value) was averaged across 4 equal sized (415.1 μm^2^) ROIs (FIJI) and assessed by unpaired t-test (GraphPad Prism v.10).

### Electrochemiluminescence quantification of amyloid beta

Flies were snap-frozen in liquid nitrogen and heads (n=25 per replicate) separated by vortexing. Heads were then homogenized for 3 mins in 150 μl of soluble extraction buffer (50 mM HEPES pH 7.3, 5 μM EDTA and 1x Protease inhibitor) on ice, vortexed, then incubated for 10 minutes at room temperature. Homogenates were sonicated at 4 °C for 4 minutes (30s on 30s off), then centrifuged at 21,000x g for 5 minutes at 4 °C. The supernatant was separated and used as the soluble fraction. 55 μl of insoluble extraction buffer (soluble extraction buffer with 5 mM Guanidinium HCL) was added to the remaining pellet. The pellet was then homogenized for 2 minutes, vortexed, and incubated for 10 minutes at room temperature. Homogenates were sonicated in a 4 °C water bath for 4 minutes (30s on 30s off), then centrifuged at 21,000x g for 5 minutes at 4°C. The supernatant was separated and used as the insoluble fraction.

The MSD V-plex assay was performed according to manufacturer’s instructions. Briefly, the plate was blocked with diluent 35 for 1 hour at room temperature. Standard calibrators were serial diluted in Diluent 35. After blocking, the plate was washed 3 times with 150 μl/well of 0.05 % Tween-PBS. Next, 25 μl of detection antibody (anti-Aβ 17-24, 4G8, mouse, MSD) was added to each well, followed by 25 µl of standard or 0.5 μg/µl brain homogenate from soluble or insoluble fractions. The plate was sealed and incubated whilst shaking for 2 hours. The plate was subsequently washed 6 times with 150 μl/well of 0.05 % Tween-PBS. The final wash was discarded and 150 μl of 2X read buffer was added to each well, before immediately reading the plate on an MSD instrument. Results were analysed via one-way ANOVAs or t-tests depending on number of groups (GraphPad Prism v.10).

### Bulk RNA sequencing

Heads of 14-day old, mated females (n=10) were dissected (N=4 reps per genotype) and RNA extracted using the RNAqueous-Micro kit (AM1931) according to the manufacturer’s instructions. Samples were sent to the Cardiff University School of Biosciences Genomics Research Hub for 6.5M reads paired-end mRNA sequencing on an Illumina NextSeq500 with a fragment length of 75 base-pairs. Quality was screened using FastQC (0.11.8); Poly-G tails, Truseq3 adapters and low-quality bases/regions (phred<3 or sliding window average of<15) were trimmed using Illumina clip (Trimmomatic (23)). Any remaining reads <20 bases in length were also removed. Reads were mapped to the *Drosophila melanogaster* genome (BDGP6.32) using STAR (24) and reads counted using featureCounts (25). Non-coding RNAs, transposons and unassigned features were removed using SARTools (26). Differential gene expression analysis was then performed using DESeq2 (27) with log_2_ fold-change shrinkage using the ashr algorithm (28). Dimensionality of variance stabilised data was assessed via principal component analyses (PCA, prcpomp). MA-plots were generated plotting normalised counts against shrunk log_2_ fold-change (threshold=±0.58) (ggplot2, RStudio v.4.4.1). Overrepresentation analysis was performed between differentially expressed genes and gene ontology molecular function (GO:MF), gene ontology biological process (GO:BP), KEGG, Wiki Pathways and TRANSFAC databases, via the functional enrichment tool g:Profiler (version e112_eg59_p19_25aa4782) (29).

### Metabolomics

Heads of 14-day old, mated females were dissected and separated into N=5 replicates of n=10 heads. Metabolites were extracted by homogenising in 1:3:1 chloroform (SLS, C2432), methanol (Fisher, 10284580) and water, followed by an hour agitation at 4 °C. MVLS Shared Research Facilities (University of Glasgow) performed Hydrophilic Interaction Liquid Chromatography (HILC) based metabolomics (UltiMate 3000 RSLC, Thermo Fisher Scientific) using a hydrophilic ZIC-pHILIC column (Merck) on samples and standards (Metabolomics Standards Initiative). Samples, were ionised using a Thermo Orbitrap QExactive (Thermo Fisher Scientific) and mass spectrometry performed in both ionisation modes, detecting metabolites in range of 70-1050 m/z. Data processing, feature annotation (based on exact mass match to compounds in databases) and identification (based on exact mass and retention time match to an authentic standard) was performed using PiMP (30). PCAs were performed on the processed data in Metaboanalyst 5.0 (31). Fragmentation analysis of the pooled samples was performed using MVLS Shared Research Facilities in-house software FrAnK. Finally, two-way ANOVAs on the top 200 features (based on variance) were used on log-transformed relative metabolite intensities using 10 % FDR in Metaboanalyst 5.0 (31,32).

## Results

### *Wwox* knockdown in neurons or glia shortens lifespan

WWOX is highly conserved between species, with *Drosophila* Wwox sharing 65% sequence similarity with human WWOX and a score of 16/16 using the DRC Integrative Ortholog Prediction Tool (DIOPT) (33). Equivalent to its mammalian orthologue (34)*, Drosophila Wwox* is expressed throughout the brain. Using a single cell transcriptomic dataset from the ageing *Drosophila* brain (17), we identified high levels of *Wwox* expression in both neuronal and glial cell-types (Fig.1A). To dissect the role of *Wwox* in both cell types, we expressed RNAi constructs under control of *elav-GAL4* (pan-neuronal) or *repo-GAL4* (pan-glial) drivers and assessed the effect on lifespan and locomotor function. *Wwox* knockdown in neurons, by two independent *UAS-RNAi* constructs (7,8,35) (Fig.S1), significantly reduced lifespan in comparison to control (RNAi Control, *Wwox^TRiP^*, *Wwox^KK^*, median lifespan=57, 42 and 40 days respectively, p_adj_<0.0001, Fig.1B). Startle-induced locomotor function, assessed by RING, was significantly decreased with age (p<0.0001) but unaffected by neuronal knockdown of *Wwox* (p=0.4458) or the interaction between the two (p=0.0778, Fig.1C). Glial knockdown of *Wwox* caused a reduction in lifespan in comparison to control (median lifespan=70 vs. 74 days, p=0.0003, Fig.1D), albeit milder than that of neuronal knockdown. Like neuronal knockdown, age significantly worsened startle-induced locomotor activity (p=0.0033), however glial knockdown of *Wwox* had no effect (p=0.0820), nor was there an interaction (p=0.5431, Fig.1E). These results suggest that both neuronal and glial *Wwox* play a role in longevity but is dispensable for startle-induced locomotor function.

**Figure 1.**
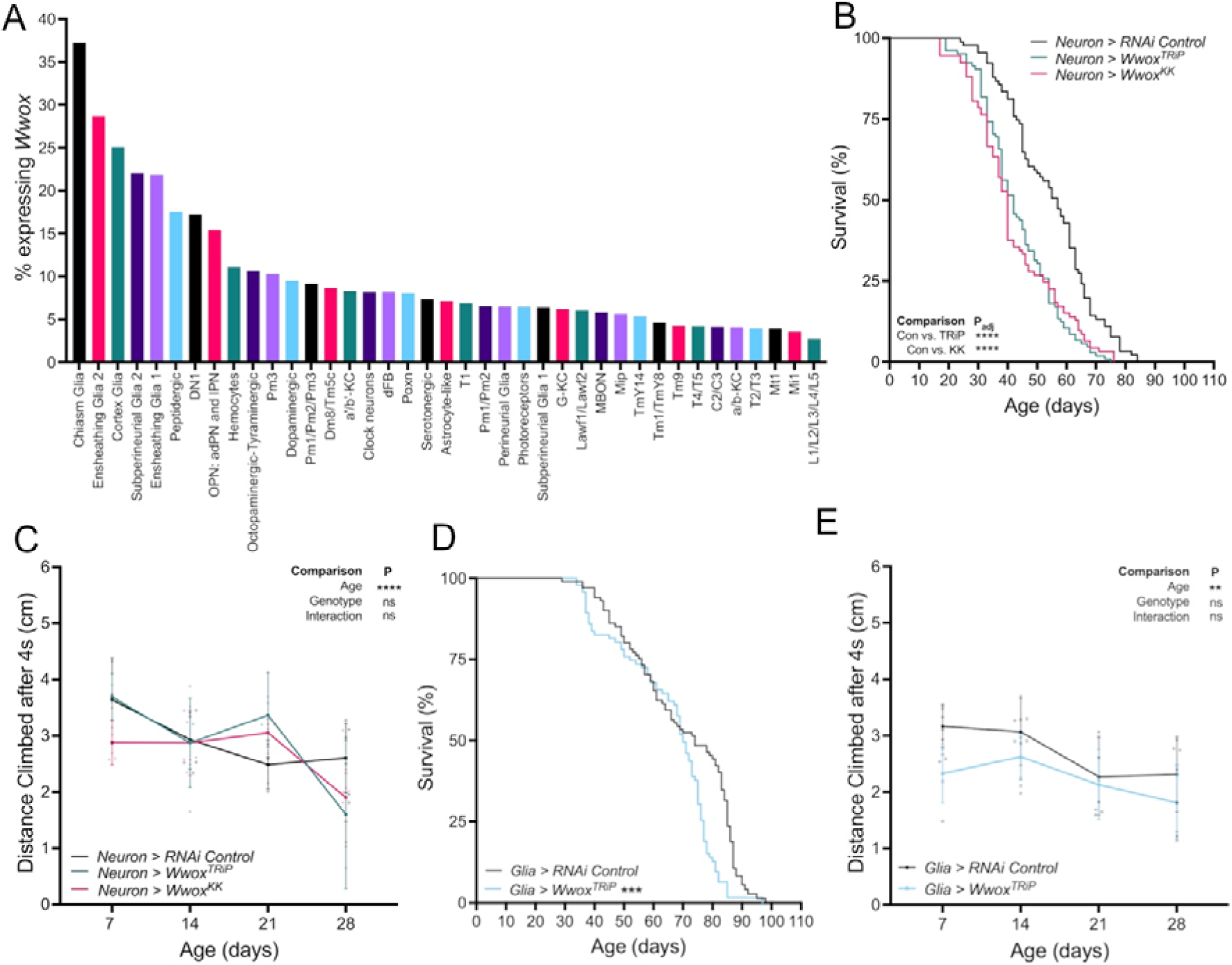
Knockdown of neuronal and glial *Wwox* reduces longevity. **A)** Analysis of single cell RNAseq data acquired through SCOPE (17) demonstrates that *Wwox* is expressed in both glia and neurons in the *Drosophila* brain. Data is presented as percentage of cells from each cluster expressing detectable levels of *Wwox* mRNA. **B)** Lifespan is reduced upon pan-neuronal *Wwox* knockdown (N=91-105 per genotype). **C)** Startle induced locomotion is unaffected by neuronal *Wwox* knockdown (N=3-6, n∼10 flies per N). **D)** Glial knockdown of *Wwox* reduces median lifespan (N=98-101). *E)* Startle induced locomotion is unaffected by glial knockdown of *Wwox* (N=4-6, n∼10 flies per N). Lifespan was assessed via (multiple) Mantel-Cox tests and locomotion via mixed effect models or two-way ANOVA with factors of age and genotype. Error bars represent standard deviation (SD), data points represent replicates. p_adj_ **<0.01, ***<0.001, ****<0.0001, ns=not significant.

### *Wwox* knockdown in neurons or glia decreases wakefulness and increases sleep

A common symptom of AD is disruption to circadian rhythms and sleep homeostasis (36–40), leading us to next to assess the contribution of neuronal or glial *Wwox* expression to sleep and circadian activity regulation. Sleep metrics were derived from activity of individually housed flies using the MATLAB toolbox SCAMP (18,19). The effect of *Wwox* manipulation was investigated on: Total activity counts, Activity counts normalised to time awake, Max wake episode duration, Mean wake episode duration, Sleep latency, Probability to transition from an active to inactive state (sleep pressure), Number of sleep episodes. Other strongly correlated sleep metrics (r>|0.8|) are shown in Table.S2.

Neuronal knockdown of *Wwox* decreased activity normalised to time awake in the light phase (p_adj_<0.0001, Fig.2A) and increased sleep pressure (p_adj_<0.0001, Fig.2B), whilst increasing mean sleep episode duration, both in the light and dark phase (both p_adj_<0.0001, Fig.2C). Similarly, pan-glial *Wwox* knockdown reduced activity normalised to time awake (p_adj_<0.0001, Fig.2D), increased sleep pressure in the light and dark phases (p_adj_<0.0001, p_adj_=0.0129, respectively, Fig.2E) and increased mean sleep episode duration (p_adj_=0.0258, Fig.2F). To understand whether these changes in sleep were due to underlying circadian differences, we evaluated the periodicity of flies in constant darkness without light entrainment cues. After neuronal knockdown of *Wwox*, all flies remained strongly rhythmic (p=0.8606, Fig.2G, I) with circadian periods comparable to control (p=0.8329, Fig.2H-I). Glial knockdown of *Wwox* had no effect on rhythmicity (p=0.2955, Fig.2J, L) or period length (p=0.3052, Fig.2K-L). *Wwox* knockdown driven by the clock specific driver *timeless-GAL4*, resulted in a similar decrease in wakefulness (Fig.S2K-R) to pan-neuronal and pan-glial knockdown, but flies remained strongly rhythmic (Fig.S2S). These results suggest that *Wwox* knockdown, irrespective of cell type, increases sleep, without disrupting circadian rhythmicity.

**Figure 2.**
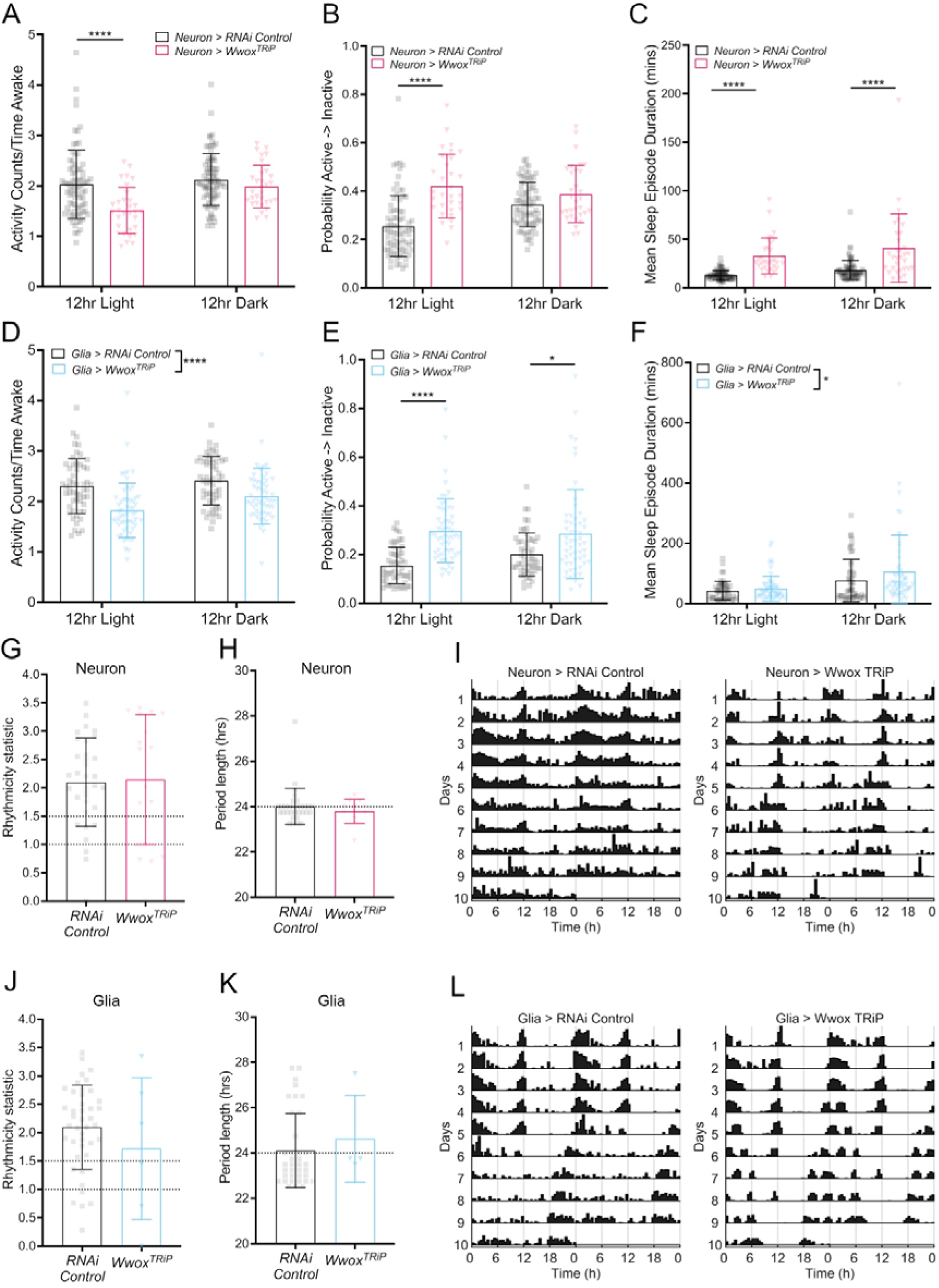
Knockdown of *Wwox* decreases wakefulness and increases sleep. Effect of pan-neuronal *Wwox* knockdown on **A)** locomotor activity, **B)** sleep pressure and **C)** mean sleep episode duration. **D)** Effect of pan-glial *Wwox* knockdown on locomotor activity, **E)** sleep pressure and **F)** mean sleep episode duration. Sleep metrics were collected over n>2 independent runs, N=31-80 individual flies per genotype. **G)** Rhythmicity statistic (RS) and **H)** period length of circadian behaviour upon pan-neuronal *Wwox* knockdown in dark (DD) conditions. **I)** Representative actograms for pan-neuronal *Wwox* knockdown. **J)** RS and **K)** period length upon pan-glial *Wwox* knockdown in DD conditions. **L)** Representative actograms for pan-glial *Wwox* knockdown. Dotted lines at RS=1 (weakly rhythmic threshold), RS=1.5 (strongly rhythmic threshold) and period length=24 hrs. Circadian rhythmicity was analysed from N=6-60 individual flies per genotype, due to removal of deaths. Whereas circadian period was assessed from N=4-57, due to the further removal of arrhythmic flies. Sleep metrics were analysed via two-way ANOVA followed by Bonferroni multiple comparisons, corrected for multiple comparisons across all sleep metrics by 5% FDR correction. p_adj_ * <0.05, ****<0.0001. Circadian data was analysed via t-test or non-parametric equivalent. Data points represent individual flies. Error bars represent SD.

### *Wwox* knockdown modifies Aβ_42_-induced phenotypes and increases Aβ_42_ load

Next, we explored the impact of *Wwox* knockdown on amyloid pathology, a hallmark feature of AD. We employed a model of amyloid toxicity whereby neuronal overexpression of aggregate-prone human Aβ_42_ harboring the Arctic mutation (E693G) and secretion signal-peptide from the *argos* gene, drives secretion of the peptide to the extracellular milieu. This causes accumulation of amyloid deposits, leading to brain vacuolation, locomotor deficits and reduction in lifespan in the fly (16). Neuronal knockdown of *Wwox* combined with overexpression of human Aβ_42_ caused a further reduction in lifespan (RNAi Control, *Wwox^TRiP^*, *Wwox^KK^*, median lifespan=41, 26 and 24 days, respectively, p_adj_<0.0001, Fig.3A). Cox-proportional hazard modelling of the effects revealed a significant interaction between Aβ_42_ overexpression and *Wwox* knockdown (p<0.0001), indicating that the decrease in lifespan is not simply the additive effect of the two variables but synergistic. As expected, startle-induced locomotor ability was significantly reduced with age (p<0.0001, Fig.3B), from ∼3.5 cm climbed at day 7 compared to only ∼1.0 cm climbed at day 28 in RNAi Control Aβ_42_ flies. There was also a significant effect of genotype (p=0.0002, Fig.3B) - multiple comparison testing revealed that both RNAi’s led to a reduction in climbing (*Wwox^TRiP^* p =0.0011 and *Wwox^KK^* p =0.0004) compared to control with knockdown flies climbing just ∼0.5 cm by day 21. We found no significant interaction between age and genotype (p=0.9323, Fig.3B). Utilising a dual-binary transgene expression system, we restricted *Wwox* knockdown to glial cells (*repo-GAL4*), whilst simultaneously restricting Aβ_42_ expression to neurons (*nSyb-QF2*). *Wwox* knockdown mildly, albeit significantly, decreased lifespan (median=22 vs. 24 days, p=0.0095, Fig.S3A) but had no effect on startle-induced locomotion (p=0.3491, Fig.S3B). These results suggest that neuronal *Wwox* may be more important than glial *Wwox* for modulation of Aβ_42_ toxicity.

**Figure 3.**
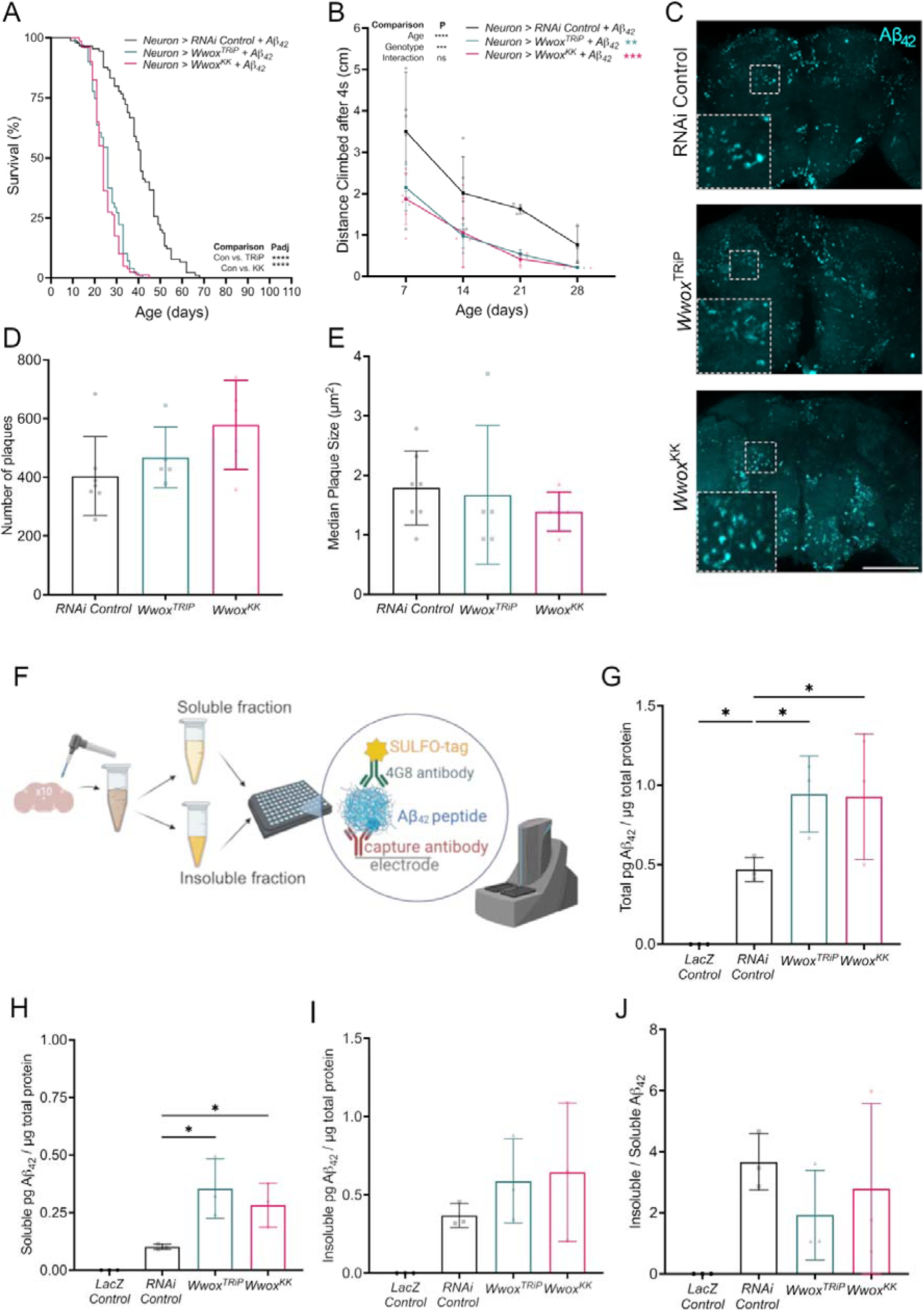
*Wwox* knockdown exacerbates Aβ_42_ related phenotypes. **A)** Lifespan is reduced upon pan-neuronal *Wwox* knockdown in the presence of Aβ_42_ overexpression (N=45-68). **B)** Startle-induced locomotion is reduced upon pan-neuronal *Wwox* knockdown in the presence of Aβ_42_ overexpression (N=2-6, n∼10 flies per N). **C)** Aβ_42_ immunocytochemistry (cyan) from day 14 *Drosophila* brains upon *Wwox* knockdown. **D)** Quantification of Aβ_42_ staining revealed no changes in number or **E)** plaque size (N=5-7 brains). **F)** Schematic of Aβ_42_ quantification in brain soluble and insoluble fractions by MSD assay. **G)** Total and **H)** soluble Aβ_42_ levels quantified by Mesoscale Discovery (MSD) assay increase with *Wwox* knockdown. **I)** But identified no changes in insoluble or **J)** the insoluble/soluble ratio (N=3, n=25 brains). Lifespan was analysed by multiple Mantel-Cox and Cox proportional hazard tests. Startle-induced locomotion was analysed by mixed effect models with factors of age and genotype. MSD assay was analysed by one-way ANOVA. p/p_adj_ *<0.05, **<0.01, ***<0.001, ****<0.0001. Error bars represent SD. Scale bar=100 µm.

To understand if neuronal *Wwox* knockdown directly impacts amyloid accumulation, we probed intact brains from these flies using the anti-Aβ [6E10] antibody. Visible Aβ_42_ deposits were quantified (Fig.3C), however no significant changes in aggregate number (p=0.1135, Fig.3D) or median size (p=0.6771, Fig.3E) were observed. To quantify changes in Aβ_42_ deposition with increased sensitivity, we assessed soluble and insoluble fractioned extracts of 14-day old fly heads by an electrochemiluminescence immunoassay (Mesoscale Discovery, MSD) using the anti-Aβ [4G8] antibody (Fig.3F). Total Aβ_42_ was significantly increased upon *Wwox* knockdown (p=0.033, Fig.3G). Interestingly, this effect was led by an increase in soluble Aβ_42_ (p=0.0024, Fig.3H), whereas insoluble fractions were only trending towards an increase (p=0.0602, Fig.3I). However, the ratio between soluble/insoluble Aβ_42_ was not significantly changed (p=0.1131, Fig.3J). Taken together, these results indicate an increase in Aβ_42_ load upon *Wwox* knockdown, without an associated increase in detectable aggregate size or number.

### Neuronal *Wwox* knockdown in the presence of Aβ_42_ alters pyruvate metabolism

To investigate the mechanisms underlying the effect of neuronal *Wwox* knockdown on Aβ_42_associated phenotypes, we performed bulk transcriptomics from flies overexpressing Aβ_42_ or a LacZ control construct, with or without neuronal *Wwox* knockdown. The effects of *Wwox* knockdown and Aβ_42_ expression could be separated along PC1 and PC2, explaining 23% and 15% of variance respectively (Fig.S4). Knockdown of *Wwox* in a control background significantly downregulated 54 genes and upregulated 44 (Fig.4A, Table.S3). Overrepresentation analysis determined these genes were enriched for oxidoreductase activity and metabolism of ascorbate, aldarate, and porphyrins (Fig.4B, Table.S3). Overexpressing human Aβ_42_ in neurons downregulated 23 genes and upregulated 51 (Fig.4C, Table.S3); overrepresentation analysis revealed a significant association of these genes with fatty acid and pyruvate metabolism (Fig.4D, Table.S3) suggesting amyloid expression induces metabolic changes. Knockdown of *Wwox* in neurons also overexpressing Aβ_42_, significantly downregulated 58 genes and upregulated 54 genes (Fig.4E, Table.S2). These genes were enriched within the pyruvate metabolism, lipid metabolism and pentose and glucuronate interconversion pathways (Fig.4F, Table.S3), suggesting that knockdown of *Wwox* in the presence of Aβ_42_ may alter the metabolome further via transcriptional regulation.

**Figure 4.**
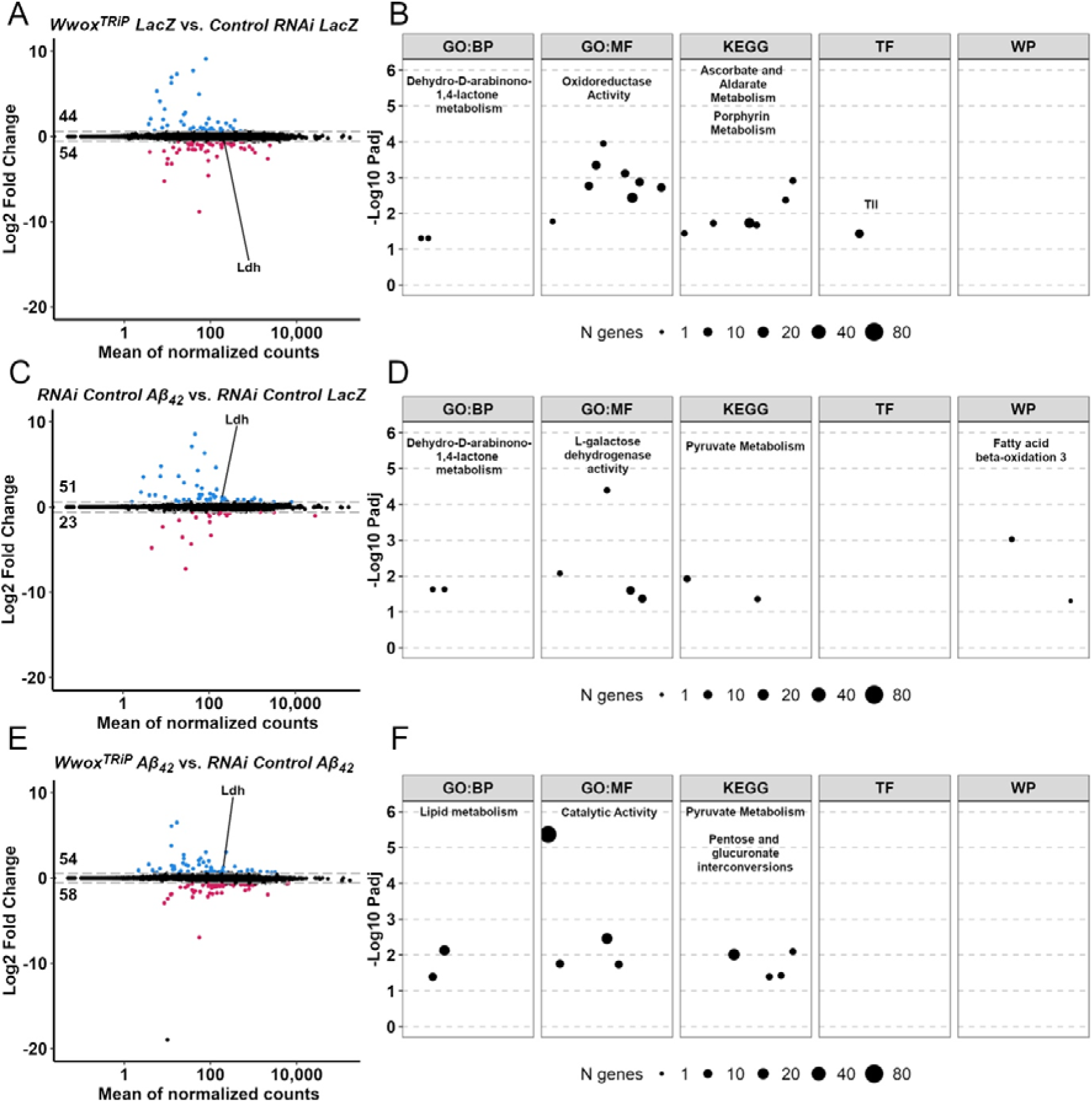
Neuronal *Wwox* knockdown in the presence of Aβ_42_ vastly alters the metabolic transcriptome. **A)** Knockdown of *Wwox* in a control background differentially expressed 98 genes, 54 down (red) and 44 up (blue). **B)** These were enriched in within metabolic pathways. **C)** MA plot summarising the significant up (51) and down regulated (23) changes in transcription after Aβ_42_ expression. **D)** These differentially expressed genes were enriched within pyruvate and fatty-acid metabolism pathways. **E)** Knockdown of *Wwox* in the presence of Aβ_42_ differentially expressed 112 genes, **F)** enriched within lipid and pyruvate metabolism terms. Dotted lines on MA plots (panels A, C and E) represent log_2_ fold-change threshold of ± 0.58. Overrepresentation analysis (panels B, D and F) was performed using g:Profiler, size of datapoints represents the number of genes identified in within the term. Abbreviations represent unique databases which were used for overrepresentation analysis GO:BP Gene ontology biological processes, GO:MF Gene ontology molecular functions, KEGG, TF TRANSFAC, WP Wiki pathways. N=4 replicates of n=10 heads per genotype.

### *Wwox* knockdown in the presence of Aβ_42_ increases lactate concentrations via Atf4

Since various metabolic pathways were identified by our transcriptomic analysis, we performed Hydrophilic Interaction Liquid Chromatography (HILC)-based metabolomics to assess the effect of overexpression of Aβ_42_ and or *Wwox* knockdown on brain metabolic profiles. Of 2,355 LC-MS peaks, 85 compounds mapped to known standards by m/z and retention time. Neuronal knockdown of *Wwox* differentially expressed eight standard-mapped metabolites, namely, reduction of L-lysine (2 peaks, both p_adj_<0.0001), orotate (p_adj_=0.0046) and 4-trimethylammoniobutanoate (p_adj_=0.0263) and upregulation of D-glucuronate (p_adj_=0.0069), 2-hydroxyglutarate (p_adj_=0.0263, 0.82 log_2_ fold-change/1.77-fold), L-lactate (p_adj_=0.0636, 0.68 log_2_ fold-change/1.60-fold) and itaconate (p_adj_=0.0885, Fig.5A, Table.S4). Aβ_42_ expression upregulated two standard-matched metabolites: 2-hydroxyglutarate (p_adj_=0.0006, 1.33 log_2_ fold-change/2.51-fold) and L-lactate (p_adj_=0.0050, 0.97 log_2_ fold-change/1.96-fold, Fig.5B, Table.S4). Both lactate and 2-hydroxyglutarate are generated from pyruvate and α-ketoglutarate, respectively, by Ldh. This is consistent with our transcriptomics analysis, which revealed a 1.19 log_2_ fold-change (2.28-fold) upregulation of *Ldh* after overexpression of Aβ_42_ (Fig.4C, Table.S3). Crucially, the interaction between neuronal *Wwox* knockdown and Aβ_42_ differentially expressed only one standard-mapped metabolite, L-lactate (p_adj_=0.0174, 1.30 log_2_ fold-change/2.46-fold Fig.5C-D, Table.S4), consistent with a 1.33 log_2_ (2.51-fold) increase in *Ldh* transcripts compared to Aβ_42_ overexpression alone (Fig.4E, Table.S3). In addition to this increase in *Ldh* and lactate, we observed corresponding decreases in the pyruvate metabolism pathway - *Pepck2*, *CG11052* and *Adh -* further supporting the role of pyruvate metabolism in the phenotypes observed (Fig.5D). An increase in *Ldh* and lactate suggests a Warburg effect-like shift towards glycolytic energy production. Interestingly however, we observed no differences in ATP levels in the heads of flies as a consequence of *Wwox* knockdown, Aβ_42_ overexpression or both combined (p=0.5720, Fig.S5A), suggesting that the observed shift towards glycolytic metabolism is sufficient to maintain energy supply and may be a protective response.

**Figure 5.**
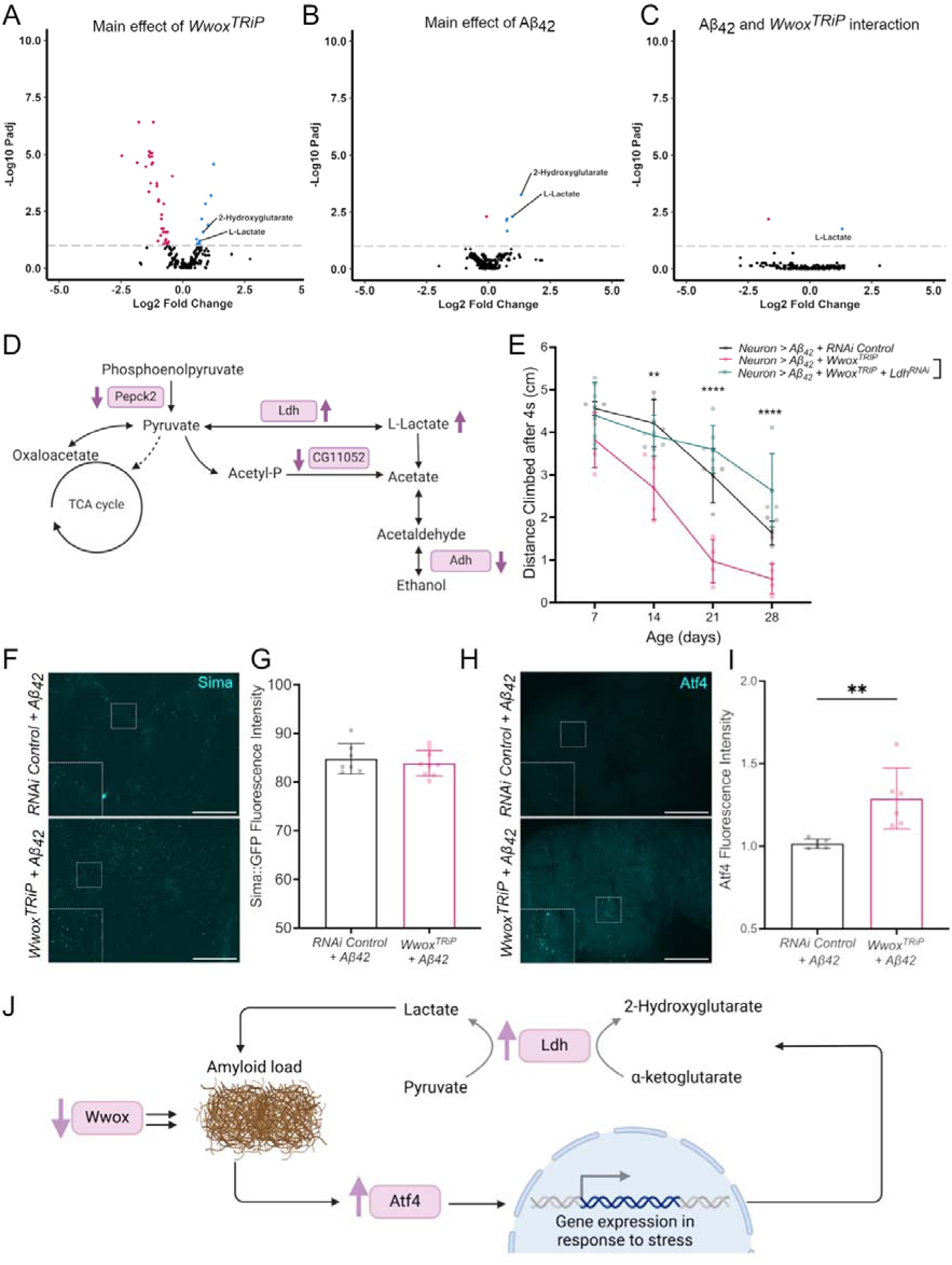
Neuronal *Wwox* knockdown in the presence of Aβ_42_ increases L-lactate levels via increased lactate dehydrogenase expression. **A)** Volcano plots summarising the significant up- and down-regulated metabolites for the main effect of *Wwox* knockdown, **B)** Aβ_42_, or **C)** the statistical interaction between the two (N=5 replicates of n=10 heads). **D)** Schematic of changes in pyruvate metabolism revealed by the statistical interaction between *Wwox* knockdown and Aβ_42_ overexpression. Differentially expressed transcripts and metabolites are depicted with arrows showing their direction of change, transcripts are represented in boxes. **E)** Knockdown of *lactate dehydrogenase* (*Ldh*) rescued startled induced locomotor deficits in *Wwox* knockdown (+Aβ_42_) flies at 14, 21 and 28 days (N=4-6 of n∼10 flies). **F)** Representative images of brains expressing the endogenously tagged Sima:GFP reporter. **G)** Quantification revealed no differences in sima levels (N=7-9 brains). **H)** ATF4 ICC in day 14 fly brains, **I)** demonstrated an upregulation of Atf4 in the brain after *Wwox* knockdown in the presence of Aβ_42_ (N=6 brains). **J)** Schematic representation of the effect of *Wwox* knockdown on pyruvate metabolism and Aβ_42_ aggregation via Atf4. Statistics=two-way ANOVA (E), t-test (G) and Mann-Whitney Test (I). p_adj_ *<0.05, **<0.01, ***<0.001, ****<0.0001. Error bars represent SD. Scale bars=100 µM. Dotted line in panels A-C represents p_adj_ threshold of 0.01.

We next investigated physiological consequences of elevated *Ldh* and lactate seen in Aβ_42_-transgenic flies with reduced *Wwox* expression. We utilized RNAi to mediate combined downregulation of both *Ldh* and *Wwox* in neurons expressing Aβ_42_ and assessed startle-induced locomotion. Multiple comparisons testing revealed that *Ldh* knockdown rescued climbing deficits seen in Aβ_42_-transgenic flies with downregulated *Wwox* (at 14-, 21- and 28-days (p_adj_=0.0081, <0.0001 and <0.0001, respectively, Fig.5E). These results suggest that *Wwox* knockdown in the presence of Aβ_42_ elevates lactate levels via neuronal *Ldh*, which is sufficient to exacerbate locomotor defects.

Both HIF1α and Atf4 are transcription factors known to promote *Ldh* expression. We investigated the levels of these proteins upon *Wwox* knockdown to further understand *Ldh* regulation in our model. No changes in endogenously tagged sima:GFP (HIF1α homolog) levels were observed in the adult fly brain after neuronal *Wwox* knockdown in Aβ_42_ overexpressing flies (p=0.5185, Fig.5F-G). However, immunocytochemistry revealed a significant upregulation of Atf4 levels in Aβ_42_ flies upon *Wwox* knockdown (p=0.0022, Fig.5H-I). Since Atf4 is activated downstream of protein kinase R (PKR)-like endoplasmic reticulum kinase (PERK) through the ISR and UPR (41), we also assessed a parallel Ire1-Xbp1 branch of the UPR, by measuring Ire1-mediated *Xbp1* splicing with a GFP-based reporter expressed in neurons (42). However, we observed no differences in neuronal spliced *Xbp1* because of *Wwox* knockdown in Aβ_42_ flies (p=0.7574, Fig.S5B-C). These data suggest that neuronal *Wwox* knockdown specifically upregulates the PERK-Atf4 branch of the ISR/UPR in Aβ_42_ flies and this could be the mechanism via which *Ldh* and lactate are upregulated, leading to exacerbated Aβ_42_-induced phenotypes (Fig.5J).

### Upregulation of neuronal *Wwox* rescues Aβ_42_ related pathology

Having described the detrimental effect of *Wwox* knockdown on Aβ_42_-induced phenotypes, we wanted to explore whether *Wwox* upregulation could also affect Aβ_42_ toxicity. We employed CRISPRa upregulation of *Wwox* by expressing UAS-dCas9-VPR and tandem gRNAs targeting the *Wwox* transcriptional start site (*Wwox^gRNA^*) or non-targeting (control^gRNA^), under pan-neuronal *elav-GAL4* control. This was sufficient to achieve ∼2-fold increase in *Wwox* mRNA levels in the fly head (Fig.S1). In the absence of Aβ_42_, *Wwox* upregulation had no significant effect on survival (p=0.1420, median=60 vs. 58 days, Fig.S6A) or startle-induced locomotion (p=0.8770, Fig.S6B). However, in the presence of neuronal Aβ_42_, upregulation of *Wwox* significantly improved lifespan (median=47.5 vs. 29 days, p<0.0001, Fig.6A) and startle-induced locomotion at days 14 and 21 (both p_adj_<0.0001, Fig.6B). We observed a similar rescue of lifespan (median=41 vs. 51 days, p=0.0021, Fig.S6C) and locomotion (p=0.0371, Fig.S6D) when overexpressing a UAS-cDNA construct of human *WWOX* (*hWWOX*) in neurons.

**Figure 6.**
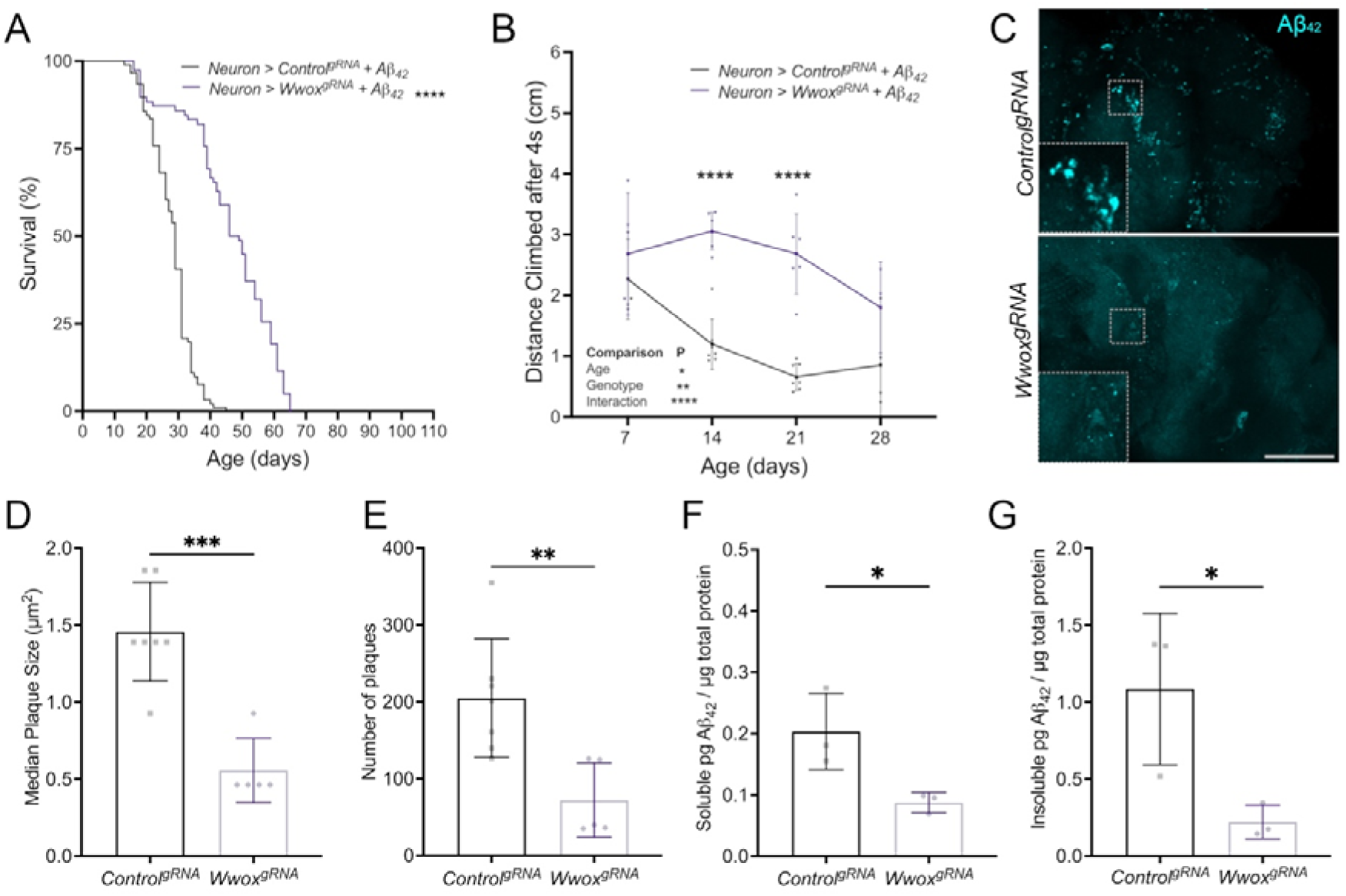
Neuronal upregulation of *Wwox* improves longevity, behaviour, and reduces amyloid load. **A)** Upregulation of *Wwox* in neurons expressing Aβ_42_ significantly improved lifespan (N=78-91), **B)** and locomotion (N=3-7 vials). **C)** Representative images of Aβ_42_ staining (cyan) in *Drosophila* brains. **D)** Quantification revealed a significant decrease in median plaque size and **E)** number (N=5-7 brains). Using the MSD assay for amyloid we revealed decreased **F)** soluble and **G)** insoluble Aβ_42_ (N=3 replicates of n=25 heads). p_adj_ *<0.05, **<0.01, ***<0.001, ****<0.0001. Error bars represent SD. Scale bar=100 µm. Statistics= Mantel-Cox tests (A), two-way ANOVA (B) and t-tests (D-G).

Overexpression of *Drosophila* or *hWWOX* cDNA in glial cells whilst simultaneously overexpressing Aβ_42_ in neurons had no effect on survival (p_adj_=0.0800 and p_adj_=0.1045 respectively, Fig.S6E) or locomotion (p=0.6654, Fig.S6F) in comparison to controls, further indicating the importance of neuronal, compared to glial *Wwox*, in the modulation of neuron-derived Aβ_42_ toxicity. In the brains of flies with *Wwox* upregulation in neurons, we found a dramatic reduction in the size (p=0.0003, Fig.6C-D) and number (p=0.0071, Fig.6C, E) of Aβ_42_ deposits with confocal microscopy. Similarly, by MSD assay, we found reduced total Aβ_42_ (p=0.0313, Fig.S6G) in both soluble (p=0.0357, Fig.6F) and insoluble (p=0.0409, Fig.6G) head lysate fractions of day 14 flies, without altering the ratio of insoluble/soluble Aβ_42_ (p=0.2530, Fig.S6H). This suggests that increasing neuronal *Wwox* can protect against Aβ_42_ associated pathology by lowering total amyloid load.

### Neuronal upregulation of *Wwox* reverses the transcriptional profile induced by Aβ_42_

To understand how neuronal upregulation of *Wwox* may provide protection against Aβ_42_ and since *Wwox* knockdown substantially altered the transcriptomic profile of Aβ_42_ flies, we again performed bulk transcriptomics on the heads of flies with *Wwox* upregulation in neurons. Upregulation of *Wwox* in a control background significantly downregulated 6 genes and upregulated none (Fig.7A, Table.S5) – these downregulated genes were enriched for fatty acid metabolism terms (Fig.7B, Table.S5). Overexpression of Aβ_42_ significantly downregulated 51 genes and upregulated 28 (Fig.7C, Table.S5), of which *Ldh* was the second most significantly upregulated gene with a 2.85 log_2_ fold-change (7.21-fold) compared to control. Overrepresentation analysis revealed dysregulation of translation repression, membrane transport and oxidative phosphorylation (Fig.7D, Table.S4). Strikingly, upregulation of neuronal *Wwox* in the presence of Aβ_42_, downregulated 305 genes and upregulated 59 (Fig.7E, Table.S4). Overrepresentation analysis revealed metabolic and immune enrichments including a dysregulation of cysteine and methionine metabolism lead largely by these downregulated genes (Fig.7F, Table.S5). To further understand these transcriptional changes, we identified 40 commonly differentially expressed genes between the analysis of Aβ_42_ vs LacZ and *Wwox^gRNA^* Aβ vs. control^gRNA^ Aβ (Table.S6). Interestingly the log_2_ fold-changes of these genes were significantly negatively correlated (r=-0.8341, p<0.0001, Fig.7G). Notably, every gene differentially expressed in Aβ_42_ vs LacZ comparison was differentially expressed in the opposing direction by the *Wwox^gRNA^* Aβ vs. control^gRNA^ Aβ_42_ comparison (Fig.7G, Table.S6). Overrepresentation analysis revealed that those genes upregulated by Aβ_42_ and downregulated by *Wwox* overexpression were enriched within immune response terms, whereas genes with the opposing change were enriched for synaptic terms (Fig.7G). Taken together, these results suggest that upregulation of *Wwox* reverses the transcriptional reprogramming of metabolic, immune and synaptic pathways caused by Aβ_42_ upregulation.

**Figure 7.**
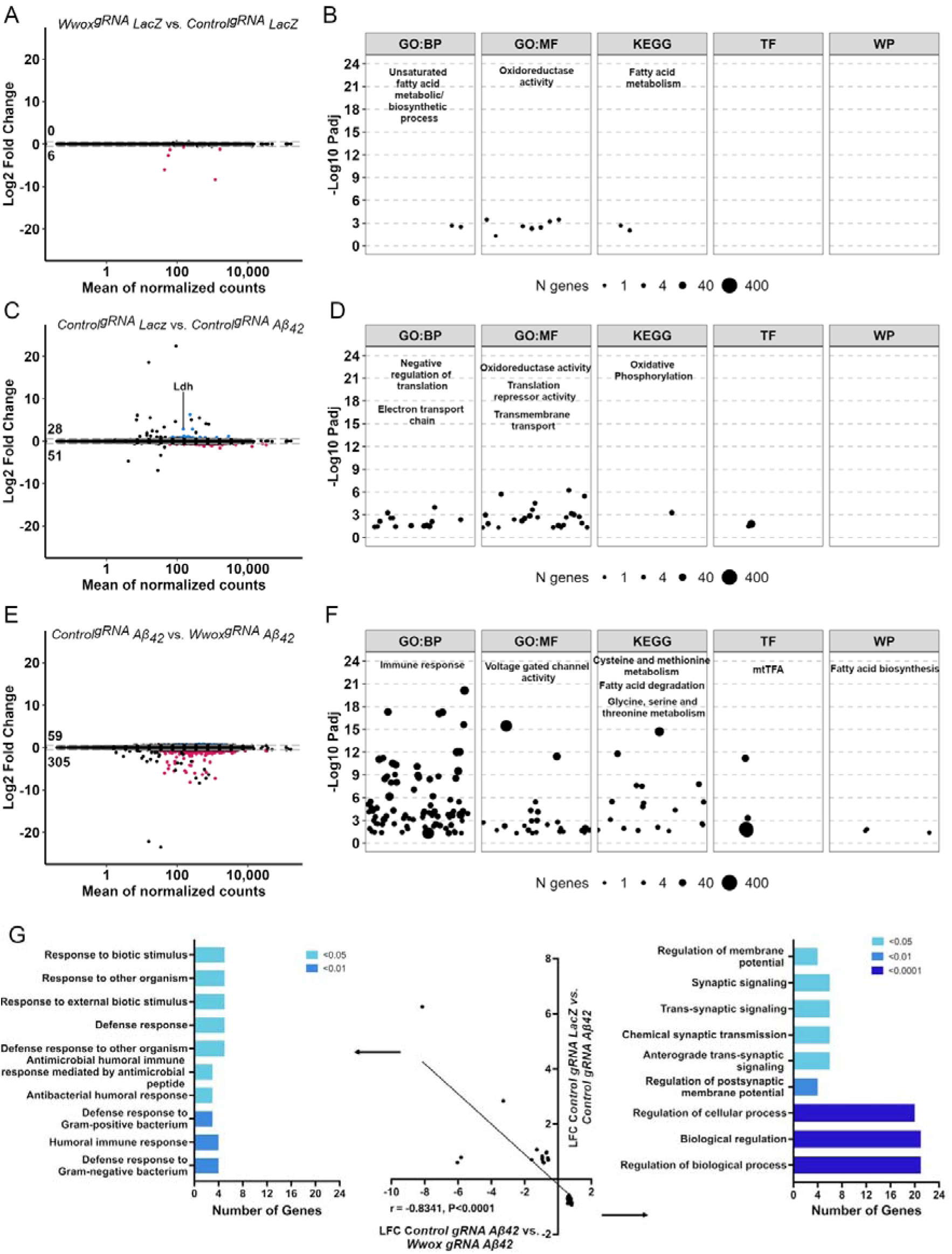
Neuronal overexpression of *Wwox* reverses the metabolic, immune and synaptic transcriptomic reprogramming caused by Aβ_42_ expression. **A)** MA plot for the effect of *Wwox* overexpression in a control background reveals dysregulation of 6 genes, **B)** g:Profiler analysis reveals enrichment within fatty acid metabolism. **C)** Overexpression of Aβ_42_ downregulated 51 genes and upregulated 28, **D)** these genes were enriched within translation repression and metabolic terms. **E)** Overexpression of *Wwox* in the presence of Aβ_42_ differentially expressed 364 genes. **F)** Overrepresentation analysis revealed many metabolic, immune and synaptic terms. Size of datapoints (panels B, D, F) represents the number of genes identified within the term. **G)** 40 genes were commonly dysregulated between Aβ_42_ expression alone and Aβ_42_ expression with *Wwox* overexpression. The log_2_ fold-change of each of these genes were significantly negatively correlated suggesting a reversal of the transcriptional profile. Those genes that were upregulated by Aβ_42_ and subsequently downregulated with *Wwox* overexpression were enriched within immune pathways (left) whereas genes that change in the opposing direction were enriched with synaptic transmission. Abbreviations represent databases used for enrichment analysis: gene ontology biological processes (GO:BP), gene ontology molecular functions (GO:MF), KEGG, TRANSFAC (TF) and Wiki Pathways (WP). Dotted lines on MA plots (panels A, C and E) represent log_2_ fold-change threshold of +/-0.58. N=4 replicates of n=10 pooled heads.

### Neuronal upregulation of *Wwox* in the presence of Aβ_42_ reduces L-methionine levels

To understand how this transcriptional reprogramming may impact the neuronal metabolism of these flies, we performed unbiased metabolomic profiling on flies expressing Aβ_42_, overexpressing *Wwox*, or overexpressing *Wwox* in the presence of Aβ_42_. Of 2414 peaks, 86 compounds were exact matches to known standards by m/z and retention time. Expression of Aβ_42_ differentially altered four standard-matched metabolites, including a reduction in sn-glycero-3-phosphocholine (p_adj_=0.0006) and an increase in betaine (p_adj_=0.0667), 2-hydroxyglutarate (p_adj_=0.0001) and citrate (p_adj_=0.0667, Fig.8A, Table.S7). This increase in 2-hydroxyglutarate is consistent with our previous finding of a 2.85 log_2_ fold-change (7.21-fold) increase in *Ldh* transcripts in Aβ_42_ expressing flies compared to control – the second most significant differentially expressed gene in this comparison (Table.S5). Neuronal *Wwox* overexpression downregulated five standard-matched metabolites – L-Leucine (p_adj_=0.0545), sn-glycero-3-phosphocholine (p_adj_=0.0667), L-methionine (p_adj_=0.0047 and 0.0091) and a suspected isomer of methyl-L-histidine (N-pi) (p_adj_=0.0545, Fig.8B, Table.S7). However, the interaction between *Wwox* overexpression and Aβ_42_ revealed just two significantly downregulated standard-matched metabolites, both representing L-methionine (p_adj_=0.0073 and p_adj_=0.0133, Fig.8C, Table.S7). Consistently, in our transcriptomics analysis we identified downregulation of seven genes within the cysteine and methionine metabolism KEGG pathway (Fig.8D). Surprisingly, the reduction of amyloid load and improvement in lifespan and locomotion of these flies therefore occurred without restoring lactate levels but instead caused a reduction in transcripts encoding numerous enzymes within the cysteine and methionine metabolism pathway, and a corresponding decrease in the metabolite L-methionine (Fig.8D-E).

**Figure 8.**
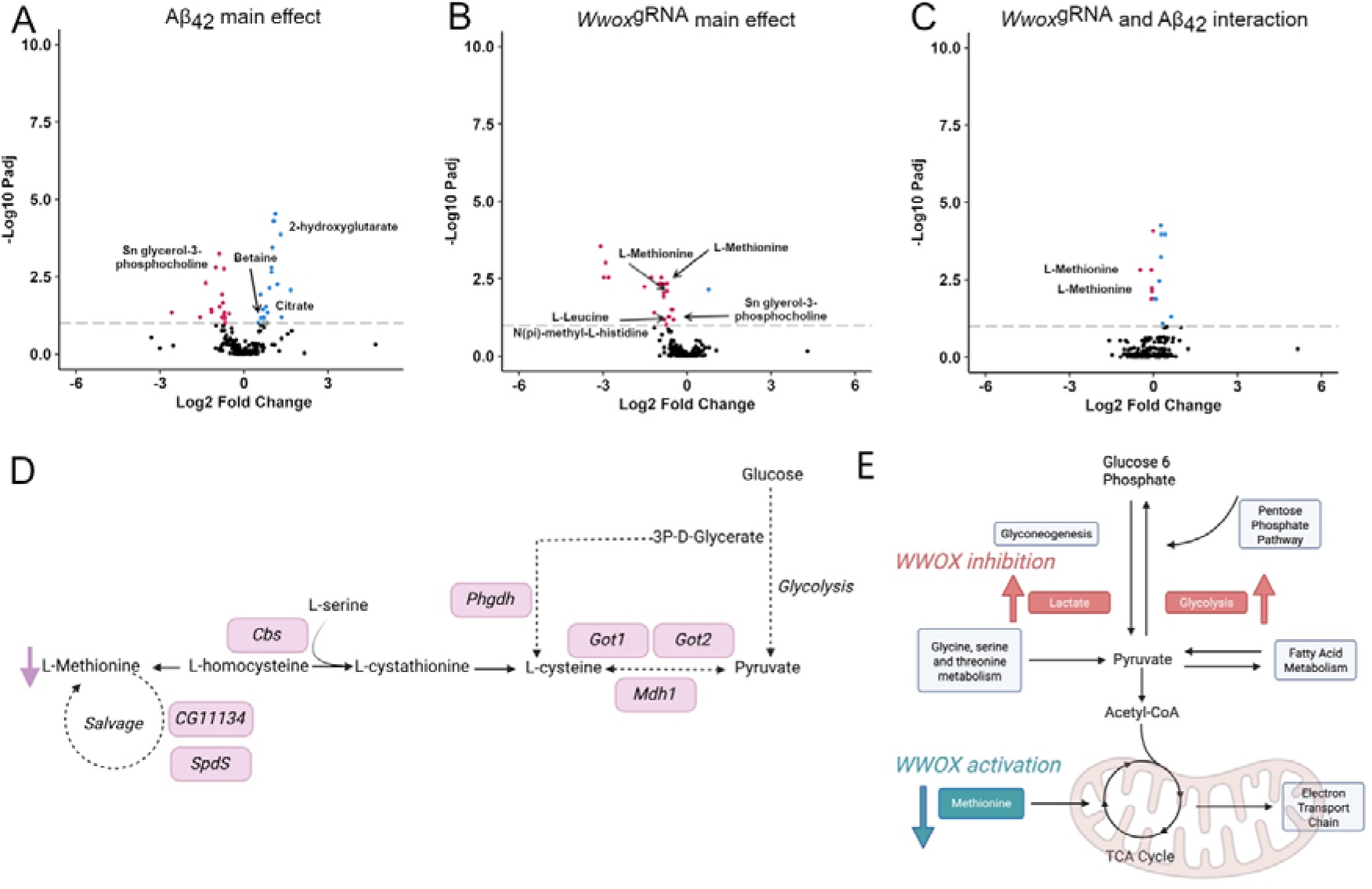
Neuronal overexpression of *Wwox* in the presence of Aβ_42_ reduces L-methionine levels. Volcano plots **A)** illustrating the differentially expressed metabolites due to Aβ_42_ expression, **B)** *Wwox* overexpression and the **C)** statistical interaction between the two. **D)** Schematic of downregulated transcripts (boxes) and metabolites (L-methionine, arrow) within the cysteine and methionine metabolism KEGG pathway revealed in our analysis of *Wwox* overexpression in the presence of Aβ_42_. **E)** Schematic to show key metabolic pathways during *Wwox* inhibition vs activation. N=5 replicates of n=10 pooled heads.

## Discussion

In this study we investigated the multifaceted, cell-type specific effects of *Wwox* manipulation in the *Drosophila* brain, in combination with the effects of ageing and human Aβ_42_ overexpression. Like mammals (34), *Drosophila Wwox* is expressed throughout the brain in both neuronal and glial cell types. Using RNAi, we reduced *Wwox* levels in these cells to recapitulate the dysregulation observed in LOAD (3,5). Whilst startle-induced locomotion was unaffected, *Wwox* knockdown in either cell type reduced lifespan, suggesting its important role in both neuronal and glial health. Additionally, *Wwox* knockdown in either cell type affected sleep-wake behaviour – decreasing activity and increasing sleep – indicating its involvement in regulating sleep homeostasis, a phenomenon also observed in early-stage LOAD (39,40). Furthermore, a study was published reporting a genome wide association between SNPs at the *WWOX* locus and self-aware sleep duration, whilst also demonstrated sleep defects in *Wwox* knockout *Drosophila*, supporting our findings (43). Our findings further show a strong interaction between sleep patterns in *Wwox* knockdown in controls animals and in our AD model, suggesting this gene could affect both age-related sleep decline and abnormalities seen in LOAD.

*Wwox* knockdown in the Aβ_42_ model of AD exacerbated neurotoxicity, leading to increased Aβ_42_ load, impaired locomotor function and reduced lifespan. However, *Wwox* knockdown in glia had no effect on the toxicity of neuron-derived Aβ_42_, suggesting that neuronal *Wwox* plays a cell-autonomous protective role against Aβ_42_-mediated toxicity. Metabolic exploration of Aβ_42_ expression revealed an increase of 2-hydroxyglutarate and L-lactate – the products of *Ldh* conversion of α-ketoglutarate and pyruvate, respectively. Consistently we found *Ldh* transcripts significantly increased with Aβ_42_ expression. This enzymatic reduction could explain previous reports of increased lactate levels in Aβ_42_-expressing *Drosophila* (13) and AD patients (44,45). The level of *Wwox* in the brain provides a link between this metabolic mechanism and genetic risk of disease. Keeping Ldh at normal levels in the brain could be key to maintaining normal metabolic homeostasis and *Ldh* overexpression has been found to shorten lifespan, disrupt circadian rhythms and induces neurodegeneration (46,47). Interestingly, *Ldh* knockdown has also been shown to cause detrimental effects in fly Aβ_42_ models, suggesting that *Ldh* expression and lactate levels need to be tightly controlled (13,48). Here we identified that *Wwox* knockdown combined with Aβ_42_ expression further elevated L-lactate levels, likely mediated by the further increase in *Ldh* expression observed in our transcriptomics data. Taken together, these data provide evidence that some genetic risk loci, like *WWOX*, can influence the metabolome to control pathology, thereby influencing disease progression.

The transcription factors Atf4 and HIF1α are both known to increase transcription of *Ldh* (11,13). While WWOX has been shown to inhibit HIF1α (6), our data suggest that Atf4, rather than the HIF1α orthologue sima, is primarily responsible for the observed *Ldh* upregulation in this fly model. Indeed, Atf4-dependent, sima-independent upregulation of *Ldh* has been previously reported in Aβ_42_-expressing *Drosophila* (13), further supporting these results. The stimulation of the PERK arm of the ISR/UPR pathway results in phosphorylation of eif2α, suppressing global translation whilst specifically promoting Atf4 translation. WWOX has previously been shown to interact in proximity with PERK, suggesting it could be directly affecting PERK activation (49). This provides a potential mechanism via which *Wwox* knockdown promotes Atf4 translation in the presence of Aβ_42_. Hippocampal neurons exposed to Aβ_42_ oligomers have been shown to increase transport of *Atf4* mRNA to axons, where it is locally translated before retrograde transport to the soma, leading to prolonged CHOP expression and CHOP-dependant cell-death (50). Whilst no *Drosophila* CHOP orthologue has been identified, prolonged PERK/Atf4 signalling has been shown to cause cell death in *Drosophila* overexpressing *Presenilin*, which encodes the catalytic subunit of Aβ-liberating γ-secretase (51). Interestingly, WWOX has also been shown to interact with GCN1 (47), an activator of Eukaryotic Translation Initiation Factor 2 Alpha Kinase 4 (GCN2) – a kinase known to phosphorylate eif2α (52). GCN2 depletion has previously been shown to be detrimental in the 5xFAD mouse model of Aβ toxicity, via enhanced PERK activation and Atf4 elevation (53). WWOX regulation of GCN2 could thus also explain the elevated Atf4 and aggravation of Aβ toxicity in our *Wwox* knockdown model.

Our findings suggest that the combined effect of *Wwox* knockdown and Aβ_42_ leads to chronic ISR/UPR activation, characterized by elevated Atf4, *Ldh* and lactate, above that of Aβ_42_ overexpression alone. Notably, we found that reducing *Ldh* levels in neurons rescued *Wwox* knockdown-induced locomotor deficits in the presence of Aβ_42,_ suggesting that elevated neuron-derived lactate levels, downstream of Atf4 signalling, could be driving the detrimental effects of *Wwox* knockdown in Aβ_42_ overexpressing flies. Lactate has been shown to be detrimental in amyloid models previously – for instance, lactic acid accumulation may increase APP-Grp78 binding, subsequent amyloidogenic cleavage and UPR disinhibition (54). Additionally, aerobic glycolysis has been suggested to directly modulate Aβ_42_ accumulation (55,56), implying a direct link between lactate and amyloid. However, further research is needed to understand the complicated dynamics between amyloid and lactate in this model of AD and how well this mirrors the human disease.

In addition to identifying deleterious effects of *Wwox* knockdown, recapitulating human risk, we found that upregulation of *Wwox* remarkably reduced amyloid load, rescued lifespan and locomotion of Aβ_42_ overexpressing flies. Surprisingly however, omics analyses show this rescue occurs without a reduction in *Ldh* or lactate, suggesting a different mechanism of protection. We observed a dramatic reversal of transcriptional changes after *Wwox* overexpression in the presence of amyloid, identifying a downregulation of metabolic and inflammatory related genes, including cysteine and methionine metabolism. Consistent with this, we observed a reduction in L-methionine as a result of this interaction. L-methionine is an essential amino acid; however, it has been suggested that diets high in methionine can result in worsened memory (57,58), neurodegeneration and increased Aβ deposition in mouse models (58). Interestingly chronic L-methionine administration has also been shown to increase inflammatory cytokine release from microglial cells (59). These results suggest that the reversal of amyloid phenotypes by *Wwox* overexpression may arise through reduction of cysteine and methionine metabolism and consequently reduction of L-methionine, inflammation and amyloid deposition.

In conclusion, our study provides new insights into the multifaceted relationship between Aβ_42_ expression and *Wwox* levels in neurons. We demonstrate that *Wwox* plays a critical role in neuronal health and the modulation of Aβ_42_ toxicity. Additionally, we identified metabolic alterations associated with Aβ_42_ expression and *Wwox* knockdown. Excitingly, increasing *Wwox* expression in neurons caused a dramatic rescue of amyloid load. Future studies should explore the potential therapeutic implications of targeting these pathways for treatment of AD.

## Declarations

The authors have no declarations.

## Data availability

Data analysis code is available at https://github.com/Fly-Cardiff. Raw reads and feature counts for the transcriptomics experiment can be accessed on Array Express, for both the knockdown (E-MTAB-14948) and overexpression (E-MTAB-14949) experiments. The metabolomics data have been deposited to MetaboLights (60) repository with the study identifier MTBLS12344.

## Contributions

GS, DM and HLC conceptualized this project and wrote the manuscript. HLC and DM performed experiments, and analysis. LA, EB and KO aided in the experiments. CR performed metabolomic experiments and analyses. OP, JH and GS were members of HLC supervisory team.

## Acknowledgements

We would like to thank the School of Biosciences Genomics Research Hub and Prof. Valentina Escott-Price at Cardiff University for their expertise regarding the transcriptomics experiments in this manuscript. We would also like to give thanks to Phil Whitfield at MVLS Shared Research Facilities (University of Glasgow) for their contribution concerning analysis of the metabolomic experiments. Schematic figures were generated with BioRender.com. This work was funded by the UKRI MRC Momentum Award (MC_PC_16030/1 to GS), the Leverhulme Trust project grant (RPG-2020-369 to GS), and the UKRI MRC Momentum Award (MC_PC_16030/2 to OP). HLC was funded by a Wellcome Trust Integrative Neuroscience PhD studentship [108891/B/15/Z]. EB and JH are supported by Alzheimer’s Research UK grant (ARUK-IRG2019B-003) and BBSRC (BB/W000865/1) awarded to JH. This work is further supported by the UK Dementia Research Institute through UK DRI Ltd, principally funded by the Medical Research Council.

## Supplementary Methods

### Quantitative polymerase chain reaction (qPCR)

12-24 hrs post eclosion, flies were snap-frozen in liquid nitrogen and heads were separated by vortexing (n=15 heads per N=3 replicate). RNA was isolated from fly heads homogenised in TRIzol reagent (Invitrogen, 15596026), according to manufacturer’s instructions. Genomic DNA contamination was removed by TURBO DNase kit (Invitrogen, AM1907) and cDNA was synthesised using the QuantiTect Reverse Transcription kit (Thermo Scientific, K0221) according to manufacturer’s instructions. cDNA was amplified using Maxima SYBR Green/ROX master mix (Invitrogen) and the following primers at 300 nM final concentration; *Wwox* forward*: ATGATAGCCCTACCCGACACA, Wwox* reverse: *TGGTTCACATAGCAAACGGTG*, *rp49* forward: *AGCATACAGGCCCAAGATCG, rp49* reverse: *TGTTGTCGATACCCTTGGGC.* The QuantStudio 7 thermocycler was used with the following conditions: Initial denaturation (95°C, 600 s), denaturation (95°C, 15 s), annealing (60 °C, 30 s), extension (72 °C, 30 s) (40 cycles), melt curve (60-95°C). Melt curve analysis was performed to ensure single-product amplification. Raw amplification data was processed using the quantitative RT PCR package in R Studio and statistical significance was calculated using a permutations approach (62).

### XBP1 imaging

We utilised the fluorescent genetically encoded reporter line *UAS-Xbp1:EGFP:HG* driven pan-neuronally to measure alternatively spliced *Xbp1*. Fresh, unfixed, brains (N=6-10) were dissected and mounted on bridge slides for immediate imaging on a Zeiss Cell Observer spinning disk confocal with an Axiocam 503 camera, using an excitation wavelength of 488 nm and 20x objective. Four regions of interest were drawn over the brain and mean gray values were calculated, avoiding trachea (FIJI, version 2.3.0). Data was analysed via t-test (GraphPad Prism v. 10) – exploring the effect of *Wwox* knockdown in the presence of amyloid.

### Celltiter-Glo 2.0

ATP concentrations were measured via the Celltiter-Glo 2.0 luciferase assay (Promega, G9242) from 14-day old flies. n=3 brains were dissected per replicate (N=4) per genotype and homogenised in 100 μl 0.1% PTX. Debris was removed via centrifugation and the supernatant was subsequently equilibrated to room temperature (30 mins) before addition of an equal volume of Celltiter-glo and a further 2 mins incubation (with rotation). After a further 10 mins luminescence was measured on a FLUostar Omega plate reader with 520 nm filter and 1.24 s exposure per well. Luminescence was averaged across four replicate wells per genotype and background luminescence subtracted. Results were then analysed via two-way ANOVA (GraphPad Prism v. 10).

### Supplementary Figure Legends

**Figure S1.**
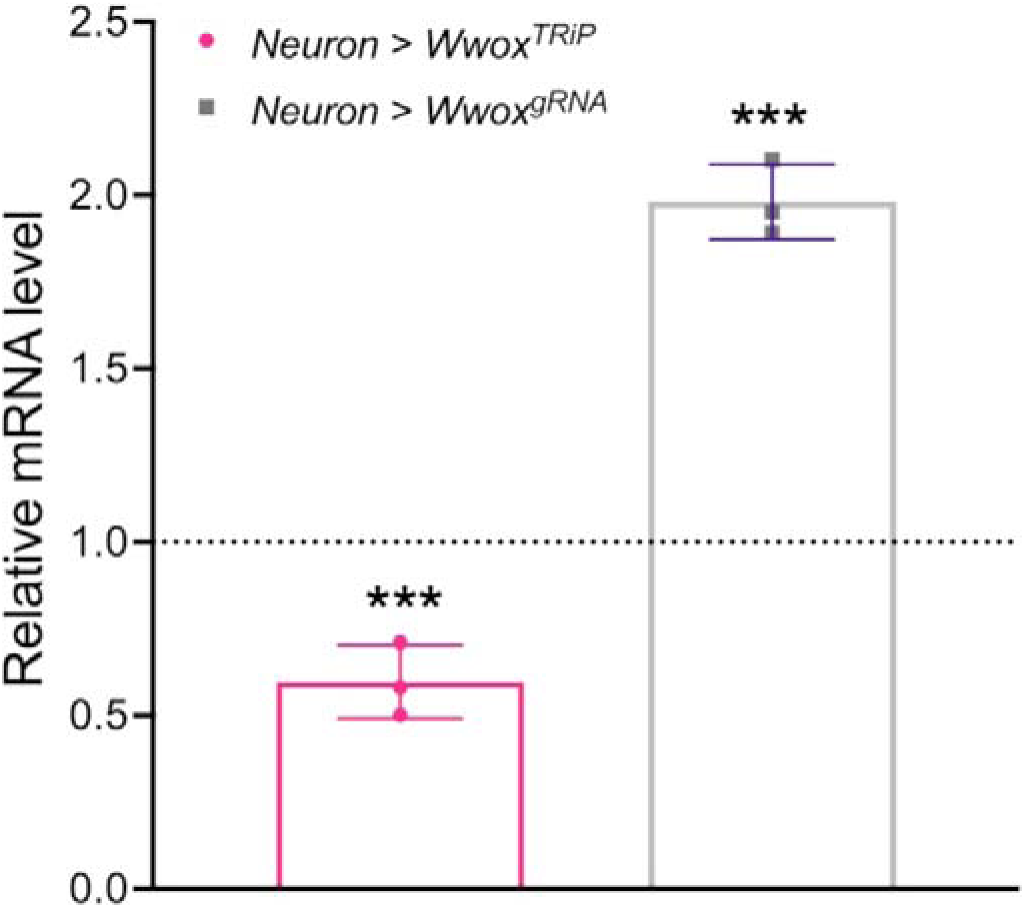
qPCR of *Wwox* mRNA levels from fly heads upon elavGAL4 driven expression of *Wwox* targeting TRiP RNAi for knockdown, or gRNA for CRISPRa upregulation. Wwox. mRNA levels were normalised to *rp49* housekeeping gene and each group was also normalised to its corresponding control genotype. mean ± SD; pairwise fixed reallocation randomization test, *** p<0.001, N=3 (15 heads per n).

**Figure S2.**
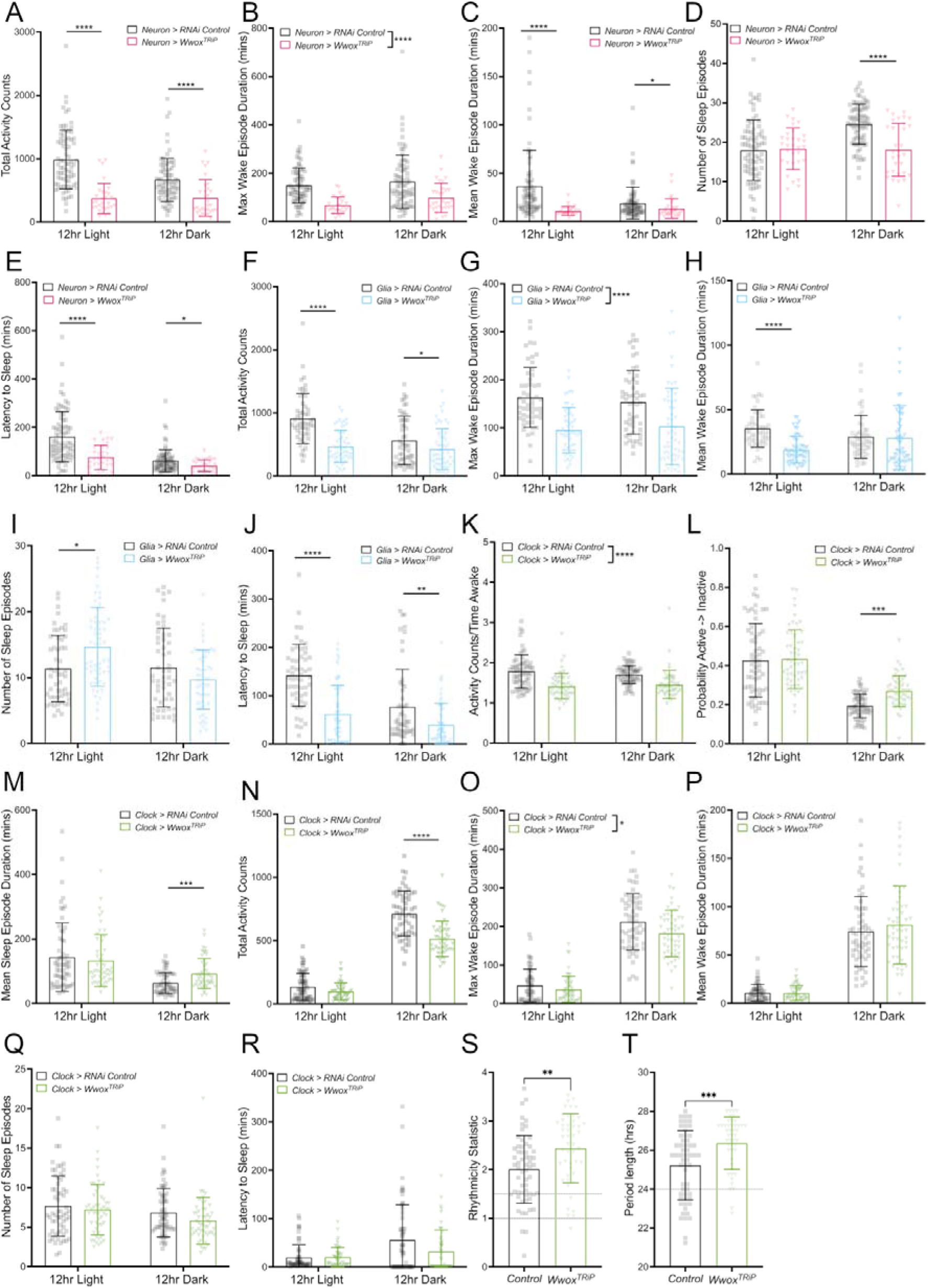
*Wwox* manipulation in neurons, glia or the clock decreases activity and increases sleep. Neuronal knockdown of *Wwox* decreases **A)** total activity counts in the light (p_adj_<0.0001) and dark (p_adj_<0.0001), **B)** maximum wake episode duration (p_adj_<0.0001) and **C)** mean wake episode duration (light p_adj_<0.0001, dark p_adj_=0.0128). **D)** The number of sleep episodes decreases (p_adj_=0.0031) **E)** as does the time taken to fall asleep (p_adj_<0.0001). Similar effects of activity are seen after glial knockdown of *Wwox*. **F)** Total activity decreases (light p_adj_<0.0001 and dark p_adj_=0.0444), **G)** max wake episode is shortened (p_adj_<0.0001), **H)** and mean wake episode is shorter during the light phase (p_adj_<0.0001). **I)** The number of sleep episodes is increased in the light phase (p_adj_=0.0107) and **J)** the latency to fall asleep is shorter (p_adj_<0.0001). Clock knockdown of *Wwox* decreases **K)** activity counts normalized to time awake (p_adj_<0.0001), **L)** increases probability to sleep (p_adj_=0.0019), and **M)** increases mean sleep episode duration (p_adj_=0.0151). Like neuronal and glial knockdown – **N)** clock knockdown decreases total activity (p_adj_<0.0001), **O)** and max wake episode duration (p_adj_=0.0403). However, no effects were seen on **P)** mean wake episode duration (p_adj_=0.5272), **Q)** number of sleep episodes (p_adj_=0.1940) or **R)** latency to sleep (p_adj_=0.4339). **S)** Circadian rhythmicity remained strong after clock knockdown of *Wwox* (p=0.0018) and **T)** circadian period was significantly lengthened (p=0.0006). two-way ANOVA followed by Bonferroni multiple comparisons were performed in A-R, P values were further corrected by 5% FDR to account for the number of metrics being assessed. Unpaired t-tests were performed in S and Mann-Whitney test in T. Dotted lines in panels S represent RS=1 (weakly rhythmic threshold), RS=1.5 (strongly rhythmic threshold) and panel T period length=24 hours. p/p_adj_ *<0.05, **<0.01, ***<0.001, ****<0.0001. Data points represent individual flies: A-E N=31-80, F-J N=53-56, K-R N=53-61, S-T N=47-60 per genotype. Error bars represent SD.

**Figure S3.**
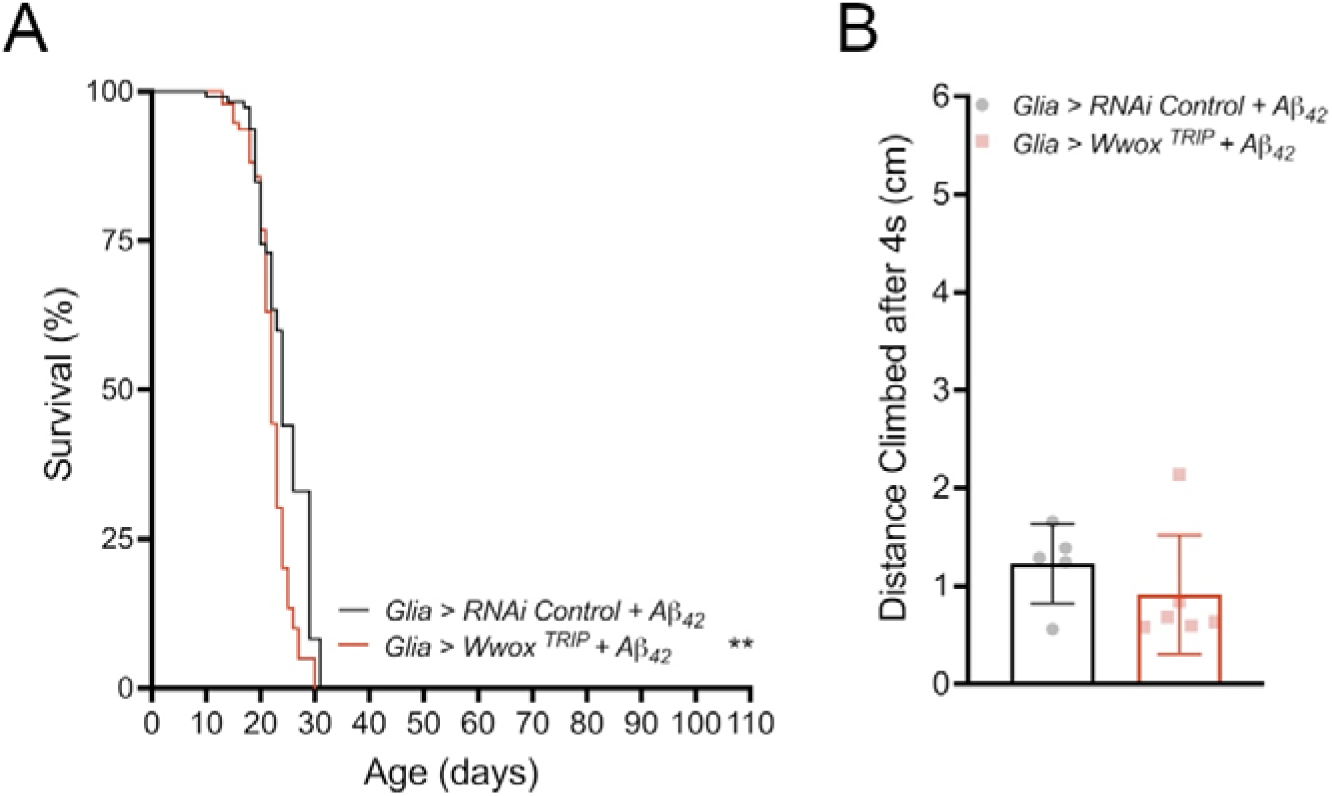
Knockdown of *Wwox* in glia whilst neurons expressed Aβ_42_ mildly effected lifespan. **A)** *Wwox* knockdown in glia, whilst expressing amyloid in neurons, decreased lifespan (p=0.0095, Mantel-Cox tests N=100-115 individual flies), **B)** but had no effect on locomotion (p= 0.3491, unpaired t-test, N=5-6 vials of n∼10). Error bars represent SD. ** p<0.01.

**Figure S4.**
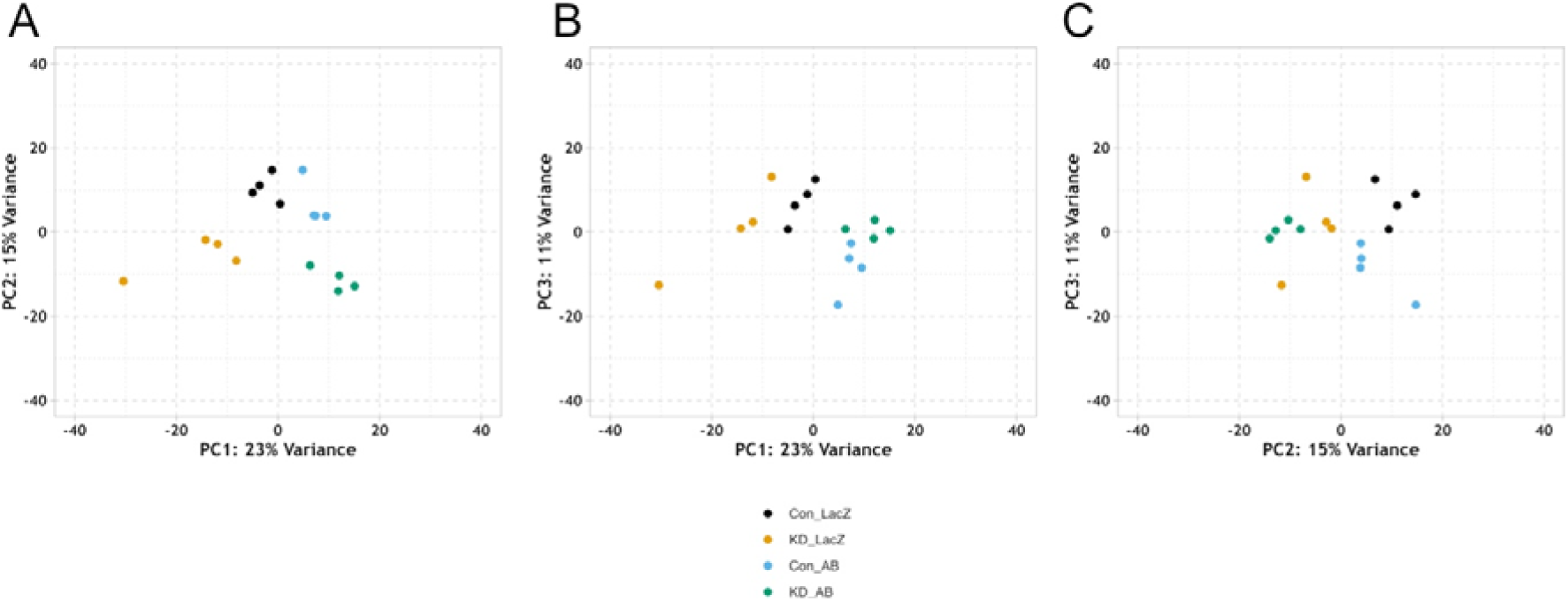
Principal component analyses show clear separation of genotypes. Genotypes from neuronal knockdown of *Wwox* transcriptomics experiment separate along **A)** PC1 (23%) vs. PC2 (15%) **B)** PC1 vs. PC3 (11%) and **C)** PC2 vs. PC3. PC1 appears to explain the effect of amyloid whereas PC2 the effect of *Wwox* knockdown. N=4 replicates from n=10 heads.

**Figure S5.**
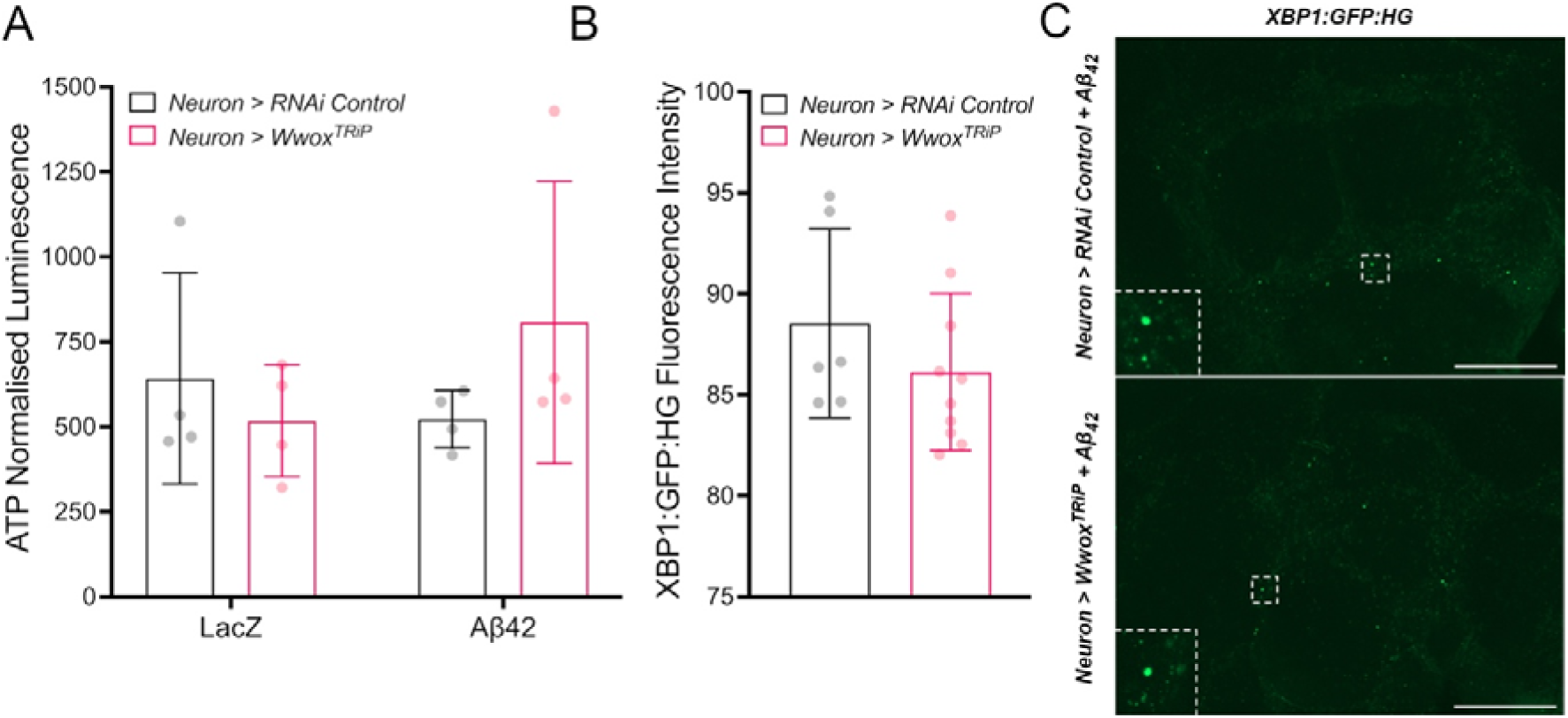
ATP levels and Xbp1-splicing is unaltered by *Wwox* knockdown in the presence of Aβ_42_. **A)** Celltitre-glo background subtracted luminescence from brain, revealed no differences in ATP concentrations after *Wwox* knockdown (p=0.5720). Mean ± SD, N=4 (n=3 brains per sample, averaged across 4 technical replicate wells). Statistics: two-way ANOVA. **B)** There are no differences in the levels of *Xbp1:GFP:HG* (reporter which only fluoresces when *Xbp1* is alternatively spliced) after *Wwox* knockdown in the presence of amyloid (p=0.2848). Error bars represent SD, statistics = t-test, N=6-10 brains. **C)** Representative images, scale bar=100 μm.

**Figure S6.**
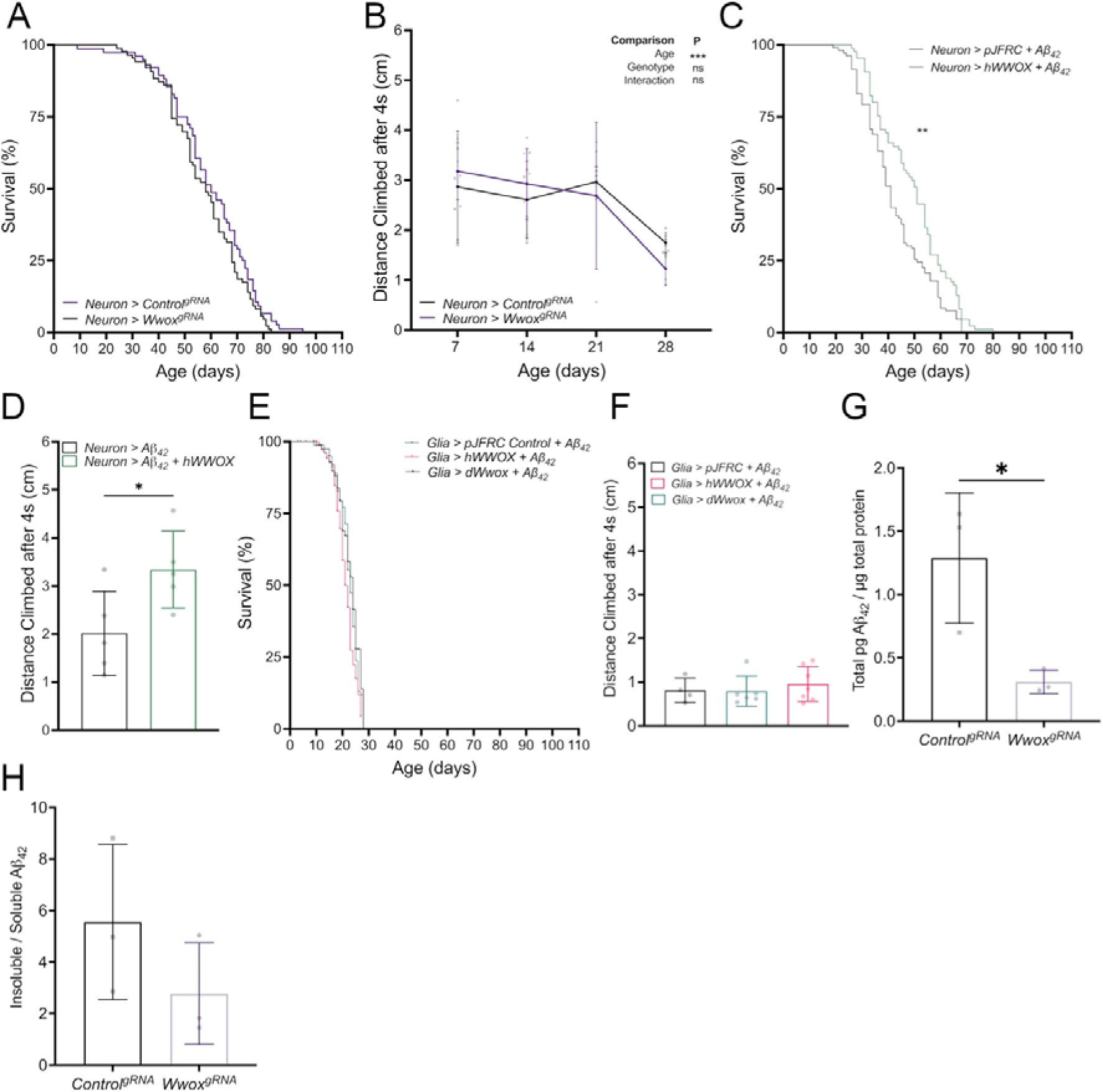
Overexpression of *Wwox* in control neurons or overexpression of *Wwox* in glia whilst expressing neuronal Aβ_42_ has no effect on behavior. **A)** There is no difference between the survival curves of flies expressing control^gRNA^ or *Wwox^gRNA^* for CRISPRa overexpression in neurons (*elav-GAL4*) in a control background (p=0.1420, N=76-86). *B)* Startle-induced locomotion of flies expressing neuronal control^gRNA^ or *Wwox^gRNA^* revealed no effect of genotype (p=0.8770, N=3-6). *C)* Expressing *hWWOX* into neurons also expressing Aβ_42_ could significantly rescue lifespan (p=0.0021, N=85-106) and *D)* improves climbing at 7-days (p=0.0371, N=5 vials). *E)* However, overexpression of *Wwox* (*Drosophila* or human) in glial cells, whilst expressing Aβ_42_ in neurons, had no effect on lifespan (p_adj_=0.8000, p_adj_=0.1045, N=84-154) or *F)* startle induced locomotion (p=0.6654, N=4-7) compared to an empty vector control. *G)* The MSD assay for amyloid revealed decreased total amyloid after *Wwox* overexpression, *H)* but no difference in the ratio of insoluble to soluble species (N=3 replicates of n=25 heads). Statistics: Mantel-Cox tests (A, C, E), two-way ANOVA (B), t-test (D, G, H), one-way ANOVA (E). Error bars represent SD. p/p_adj_ * <0.05, **<0.01, ***<0.001, ns=non-significant.

**Figure S7.**
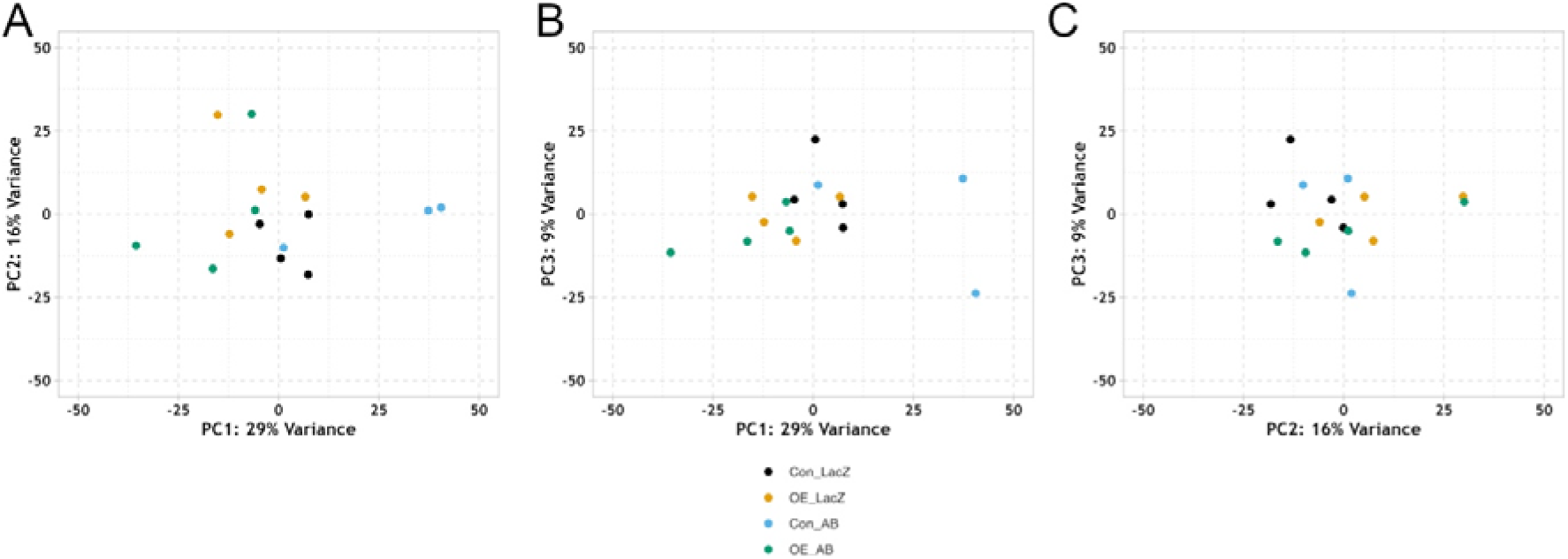
Principal component analyses for overexpression transcriptomics. Principal component analysis reveals that 29% of the variation in the dataset can be explained by PC1, 16% by PC2 and 9% by PC3. **A)** Comparison of PC1 vs PC2 reveals separation of control and Wwox overexpression along PC1. There is less clear separation between the genotypes when comparing **B)** PC1 and PC2, or **C)** PC2 and PC3. N=4 replicates of n=10 heads.

### Supplementary Tables

Uploaded in a separate file with this manuscript.

## References

1. Bellenguez C, Küçükali F, Jansen IE, Kleineidam L, Moreno-Grau S, Amin N, et al. New insights into the genetic etiology of Alzheimer’s disease and related dementias. Nat Genet. 2022;54(4):412–36.

2. Kunkle BW, Grenier-Boley B, Sims R, Bis JC, Damotte V, Naj AC, et al. Genetic meta-analysis of diagnosed Alzheimer’s disease identifies new risk loci and implicates Aβ, tau, immunity and lipid processing. Nat Genet. 2019;51(3):414–30.

3. Patel D, Zhang X, Farrell JJ, Chung J, Stein TD, Lunetta KL, et al. Cell-type Specific Expression Quantitative Trait Loci Associated with Alzheimer Disease in Blood and Brain Tissue. medRxiv. 2020;(617).

4. Stevenson-Hoare J, Heslegrave A, Leonenko G, Fathalla D, Bellou E, Luckcuck L, et al. Plasma biomarkers and genetics in the diagnosis and prediction of Alzheimer’s disease. Brain [Internet]. 2023;146(2):690–9. Available from: 10.1093/brain/awac128

5. Sze CI, Su M, Pugazhenthi S, Jambal P, Hsu LJ, Heath J, et al. Down-regulation of WW domain-containing oxidoreductase induces Tau phosphorylation in vitro: A potential role in Alzheimer’s disease. Journal of Biological Chemistry. 2004;279(29):30498–506.

6. Abu-Remaileh M, Seewaldt VL, Aqeilan RI. WWOX loss activates aerobic glycolysis. Mol Cell Oncol. 2015;2(2).

7. O’Keefe L V., Colella A, Dayan S, Chen Q, Choo A, Jacob R, et al. Drosophila orthologue of WWOX, the chromosomal fragile site FRA16D tumour suppressor gene, functions in aerobic metabolism and regulates reactive oxygen species. Hum Mol Genet. 2011;20(3):497–509.

8. Choo A, O’Keefe L V., Lee CS, Gregory, Stephen L, Shaukat Z, Colella A, et al. Tumor Suppressor WWOX Moderates the Mitochondrial Respiratory Complex. Genes Chromosomes Cancer. 2015;54:745–61.

9. Bednarek AK, Keck-Waggoner CL, Daniel RL, Laflin KJ, Bergsagel PL, Kiguchi K, et al. WWOX, the FRA16D gene, behaves as a suppressor of tumor growth. Cancer Res. 2001;61(22):8068–73.

10. Bednarek AK, Laflin KJ, Daniel RL, Liao Q, Hawkins KA, Aldaz CM. WWOX, a novel WW domain-containing protein mapping to human chromosome 16q23.3-24.1, a region frequently affected in breast cancer. Cancer Res. 2000;60(8):2140–5.

11. Lee JW, Bae SH, Jeong JW, Kim SHSHK, Kim KW. Hypoxia-inducible factor (HIF-1)a: its protein stability and biological functions. Exp Mol Med [Internet]. 2004;36(1):1–12. Available from: https://www.nature.com/articles/emm20041

12. Abu-Remaileh M, Aqeilan RI. Tumor suppressor WWOX regulates glucose metabolism via HIF1α modulation. Cell Death Differ. 2014;21(11):1805–14.

13. Niccoli T, Kerr F, Snoeren I, Fabian D, Aleyakpo B, Ivanov D, et al. Activating transcription factor 4-dependent lactate dehydrogenase activation as a protective response to amyloid beta toxicity. Brain Commun. 2021;3(2):1–13.

14. Mertens J, Herdy JR, Traxler L, Schafer ST, Schlachetzki JCM, Böhnke L, et al. Age-dependent instability of mature neuronal fate in induced neurons from Alzheimer’s patients. Cell Stem Cell. 2021;28(9):1533–1548.e6.

15. Traxler L, Herdy JR, Stefanoni D, Eichhorner S, Pelucchi S, Szücs A, et al. Warburg-like metabolic transformation underlies neuronal degeneration in sporadic Alzheimer’s disease. Cell Metab. 2022;34(9):1248–1263.e6.

16. Crowther DC, Kinghorn KJ, Miranda E, Page R, Curry JA, Duthie FAI, et al. Intraneuronal Aβ, non-amyloid aggregates and neurodegeneration in a Drosophila model of Alzheimer’s disease. Neuroscience. 2005;132(1):123–35.

17. Davie K, Janssens J, Koldere D, De Waegeneer M, Pech U, Kreft Ł, et al. A Single-Cell Transcriptome Atlas of the Aging Drosophila Brain. Cell. 2018;174(4):982–998.e20.

18. Donelson N, Kim EZ, Slawson JB, Vecsey CG, Huber R, Griffith LC. High-resolution positional tracking for long-term analysis of Drosophila sleep and locomotion using the “tracker” program. PLoS One. 2012;7(5).

19. Vecsey CG, Koochagian C, Porter MT, Roman G, Sitaraman D. Analysis of Sleep and Circadian Rhythms from Drosophila Activity-Monitoring Data Using SCAMP. Cold Spring Harb Protoc. 2024 Nov 9;

20. Hendricks JC, Finn SM, Panckeri KA, Chavkin J, Williams JA, Sehgal A, et al. Rest in Drosophila is a sleep-like state. Neuron. 2000;25:129–38.

21. Shaw PJ, Cirelli C, Greenspan RJ, Tononi G. Correlates of sleep and waking in Drosophila melanogaster. Science (1979). 2000;287(5459):1834–7.

22. Hunt RJ, Granat L, McElroy GS, Ranganathan R, Chandel NS, Bateman JM. Mitochondrial stress causes neuronal dysfunction via an ATF4-dependent increase in L-2-hydroxyglutarate. Journal of Cell Biology. 2019;218(12):4007–16.

23. Bolger AM, Lohse M, Usadel B. Trimmomatic: A flexible trimmer for Illumina Sequence Data. Bioinformatics,. 2014;btu170.

24. Dobin A, Davis CA, Schlesinger F, Drenkow J, Zaleski C, Jha S, et al. STAR: Ultrafast universal RNA-seq aligner. Bioinformatics. 2013;29(1):15–21.

25. Liao Y, Smyth GK, Shi W. FeatureCounts: An efficient general purpose program for assigning sequence reads to genomic features. Bioinformatics. 2014;30(7):923–30.

26. Varet H, Brillet-Guéguen L, Coppée JY, Dillies MA. SARTools: A DESeq2- and edgeR-based R pipeline for comprehensive differential analysis of RNA-Seq data. PLoS One. 2016;11(6):1–8.

27. Love MI, Huber W, Anders S. Moderated estimation of fold change and dispersion for RNA-seq data with DESeq2. Genome Biol. 2014;15(12):550.

28. Stephens M. False discovery rates: A new deal. Biostatistics. 2017 Apr 1;18(2):275–94.

29. Raudvere U, Kolberg L, Kuzmin I, Arak T, Adler P, Peterson H, et al. G:Profiler: A web server for functional enrichment analysis and conversions of gene lists (2019 update). Nucleic Acids Res. 2019;47(W1):W191–8.

30. Gloaguen Y, Morton F, Daly R, Gurden R, Rogers S, Wandy J, et al. PiMP my metabolome: An integrated, web-based tool for LC-MS metabolomics data. Bioinformatics. 2017;33(24):4007–9.

31. Pang Z, Zhou G, Ewald J, Chang L, Hacariz O, Basu N, et al. Using MetaboAnalyst 5.0 for LC-HRMS spectra processing, multi-omics integration and covariate adjustment of global metabolomics data. Nat Protoc. 2022;

32. Creek DJ, Jankevics A, Burgess K, Breitling R, Barrett MP. IDEOM: an Excel interface for analysis of LC–MS-based metabolomics data. Bioinformatics. 2012;28(7):1048–9.

33. Hu Y, Flockhart I, Vinayagam A, Bergwitz C, Berger B, Perrimon N, et al. An integrative approach to ortholog prediction for disease-focused and other functional studies. BMC Bioinformatics. 2011;12.

34. Marcelo Aldaz C, Hussain T. Wwox loss of function in neurodevelopmental and neurodegenerative disorders. Int J Mol Sci. 2020;21(23):1–22.

35. O’Keefe L V., Lee CS, Choo A, Richards RI. Tumor suppressor WWOX contributes to the elimination of tumorigenic cells in drosophila melanogaster. PLoS One. 2015;10(8):1–19.

36. Wadsworth LP, Lorius N, Donovan NJ, Locascio JJ, Rentz DM, Johnson KA, et al. Neuropsychiatric symptoms and global functional impairment along the Alzheimer’s continuum. Dement Geriatr Cogn Disord. 2012;34(2):96–111.

37. Van Someren EJW, Hagebeuk EEO, Lijzenga C, Scheltens P, De Rooij SEJA, Jonker C, et al. Circadian rest-activity rhythm disturbances in Alzheimer’s disease. Biol Psychiatry. 1996;40(4):259–70.

38. Coogan AN, Schutová B, Husung S, Furczyk K, Baune BT, Kropp P, et al. The circadian system in Alzheimer’s disease: Disturbances, mechanisms, and opportunities. Biol Psychiatry. 2013;74(5):333–9.

39. Ju YES, Lucey BP, Holtzman DM. Sleep and Alzheimer disease pathology-a bidirectional relationship. Nat Rev Neurol. 2014;10(2):115–9.

40. Ju YES, McLeland JS, Toedebusch CD, Xiong C, Fagan AM, Duntley SP, et al. Sleep quality and preclinical Alzheimer disease. JAMA Neurol. 2013;70(5):587–93.

41. Hetz C, Mollereau B. Disturbance of endoplasmic reticulum proteostasis in neurodegenerative diseases. Nat Rev Neurosci. 2014;15(4):233–49.

42. Souid S, Lepesant JA, Yanicostas C. The xbp-1 gene is essential for development in Drosophila. Dev Genes Evol. 2007 Feb;217(2):159–67.

43. Kim S, Kang SW, Kim SE, Kim HJ, Kim SA, Lee YW, et al. Genome-wide identification and functional validation of the WW domain containing oxidoreductase gene associated with sleep duration. Sci Rep. 2025 Dec 1;15(1):5552.

44. Liguori C, Chiaravalloti A, Sancesario G, Stefani A, Sancesario GM, Mercuri NB, et al. Cerebrospinal fluid lactate levels and brain [18F]FDG PET hypometabolism within the default mode network in Alzheimer’s disease. Eur J Nucl Med Mol Imaging [Internet]. 2016;43(11):2040–9. Available from: 10.1007/s00259-016-3417-2

45. Liguori C, Stefani A, Sancesario G, Sancesario GM, Marciani MG, Pierantozzi M. CSF lactate levels, τ proteins, cognitive decline: A dynamic relationship in Alzheimer’s disease. J Neurol Neurosurg Psychiatry. 2015;86(6):655–9.

46. Long DM, Frame A, Reardon PN, Cumming RC, Hendrix DA, Kretzschmar D, et al. Lactate dehydrogenase expression modulates longevity and neurodegeneration in Drosophila melanogaster. Aging [Internet]. 2020;12(11). Available from: www.aging-us.com

47. Hussain T, Lee J, Abba MC, Chen J, Marcelo Aldaz C. Delineating WWOX protein interactome by tandem affinity purification-mass spectrometry: Identification of top interactors and key metabolic pathways involved. Front Oncol. 2018;8(DEC):1–14.

48. Frame AK, Robinson JW, Mahmoudzadeh NH, Tennessen JM, Simon AF, Cumming RC. Aging and memory are altered by genetically manipulating lactate dehydrogenase in the neurons or glia of flies. Aging. 2023;15(4):947–81.

49. Sassano ML, Derua R, Waelkens E, Agostinis P, van Vliet AR. Interactome Analysis of the ER Stress Sensor Perk Uncovers Key Components of ER-Mitochondria Contact Sites and Ca2+ Signalling. Contact. 2021 Jan 1;4.

50. Baleriola J, Walker CA, Jean YY, Crary JF, Troy CM, Nagy PL, et al. Axonally synthesized ATF4 transmits a neurodegenerative signal across brain regions. Cell. 2014 Aug 28;158(5):1159–72.

51. Demay Y, Perochon J, Szuplewski S, Mignotte B, Gaumer S. The PERK pathway independently triggers apoptosis and a Rac1/Slpr/JNK/Dilp8 signaling favoring tissue homeostasis in a chronic ER stress Drosophila model. Cell Death Dis. 2014 Jan 1;5(10).

52. Vazquez De Aldana CR, Maiton MJ, Hinnebusch AG. GCN20, a novel ATP binding cassette protein, and GCN1 reside in a complex that mediates activation of the elF-2α kinase GCN2 in amino acid-starved cells. EMBO Journal. 1995;14(13):3184–99.

53. Devi L, Ohno M. Deletion of the eIF2α Kinase GCN2 Fails to Rescue the Memory Decline Associated with Alzheimer’s Disease. PLoS One. 2013 Oct 11;8(10).

54. Xiang Y, Xu G, Weigel-Van Aken AK. Lactic acid induces aberrant amyloid precursor protein processing by promoting its interaction with endoplasmic reticulum chaperone proteins. PLoS One. 2010;5(11):1–8.

55. Vlassenko AG, Vaishnavi SN, Couture L, Sacco D, Shannon BJ, Mach RH, et al. Spatial correlation between brain aerobic glycolysis and amyloid-β (Aβ) deposition. Proc Natl Acad Sci U S A. 2010;107(41):17763–7.

56. Bero AW, Yan P, Roh JH, Cirrito JR, Stewart FR, Raichle ME, et al. Neuronal activity regulates the regional vulnerability to amyloid-β 2 deposition. Nat Neurosci. 2011;14(6):750–6.

57. Kalani A, Chaturvedi P, Kalani K, Kamat PK, Chaturvedi P, Tyagi N. A high methionine, low folate and Vitamin B6/B12 containing diet can be associated with memory loss by epigenetic silencing of netrin-1. Neural Regen Res. 2019 Jul 1;14(7):1247–54.

58. Pi T, Wei S, Jiang Y, Shi JS. High Methionine Diet-Induced Alzheimer’s Disease like Symptoms Are Accompanied by 5-Methylcytosine Elevated Levels in the Brain. Behavioural Neurology. 2021;2021.

59. Alachkar A, Agrawal S, Baboldashtian M, Nuseir K, Salazar J, Agrawal A. L-methionine enhances neuroinflammation and impairs neurogenesis: Implication for Alzheimer’s disease. J Neuroimmunol. 2022 May 15;366.

60. Yurekten O, Payne T, Tejera N, Amaladoss FX, Martin C, Williams M, et al. MetaboLights: open data repository for metabolomics. Nucleic Acids Res. 2024 Jan 5;52(D1):D640–6.

61. Kaneko M, Hall JC. Neuroanatomy of cells expressing clock genes in Drosophila: Transgenic manipulation of the period and timeless genes to mark the perikarya of circadian pacemaker neurons and their projections. Journal of Comparative Neurology. 2000;422(1):66–94.

62. Ritz C, Spiess AN. qpcR: An R package for sigmoidal model selection in quantitative real-time polymerase chain reaction analysis. Bioinformatics. 2008;24(13):1549–51.

